# Metformin decreases *Cyp26a1* to prevent hepatocarcinogenesis through down- regulating CD8^+^ T cells

**DOI:** 10.1101/2022.04.27.489721

**Authors:** Weizhi He, Miaomiao Chen, Chong Li, Xicheng Wang, Wenjian Chen, Lili Pan, Yangyang Cui, Zhao Yu, Guoxiu Wu, Yang Yang, Qinghe Tang, Jinghan Wang, Zhiying He

## Abstract

Hepatocellular carcinoma (HCC) is a highly heterogeneous cancer, which limits the selectivity of prevention and treatment. Preclinical and clinical studies suggested that in patients with diabetes, prolonged use of metformin, the AMPK activator, was associated with a reduction of HCC incidence. This association promotes us to investigate the possible functions and mechanisms of metformin in HCC without diabetes backgrounds. Here, we found that several unique pathways that changed during chronic liver injury of *Fah^-/-^* mice, including glucose metabolic process and retinol metabolism. Further, metformin suppressed the tumor formation in chronic liver injury of *Fah^-/-^* mice. RNA sequencing, *in vivo* and *in vitro* experiments showed that metformin suppressed *Cyp26a1* gene expression of hepatocyte. Moreover, the down-regulation of Cyp26a1 leads to the increased level of all-trans-retinoic acid (atRA), which could suppress the tumor formation in our model. On the other hand, flow multicolor analysis showed that the cell number and proportion of cancer promoting (pro-tumor) CD8^+^ T cells increased significantly during chronic liver injury in *Fah^-/-^* mice, and both metformin and atRA treatment could reduce the number and proportion of pro-tumor CD8^+^ T cells. We also found metformin decreased the *Cyp26a1* expression through the AMPK/JNK/c-Jun pathway. In short, the association between the metformin and atRA may explain the commonness of their anti-tumor activities. Our findings highlight the importance of targeting the precancerous microenvironment for the prognosis, prevention and treatment of HCC.

## Introduction

Hepatocellular carcinoma (HCC) is the sixth most common neoplasm and the third leading cause of cancer death in the world (1). Although HCC treatment has made great advances over the past decades, tumor heterogeneity of the HCC still keeps its mortality rate high. HCC is a complex malignancy that can be triggered by various factors, including hepatic viral infections, environmental exposure to toxic substance (such as aflatoxin B1), alcohol abuse and non-alcoholic fatty liver disease (NAFLD) (2). Besides, growing evidence supports an association between HCC and metabolic syndrome, diabetes or obesity (3). In addition to these acquired HCC, patients with certain gene defects are bound to develop HCC (4-6). The heterogeneity of HCC, especially the diverseness in causes and progressions, complicates its therapeutic approach.

Metformin (Met), a synthetic analog of guanidine, is the most commonly prescribed drug in treating the type 2 diabetes mellitus (T2DM) (7). In addition to lowering glucose, metformin directly inhibits complex I (CI) of the electron transport chain, resulting in decreased complex I activity and oxidative phosphorylation (OXPHOS) level in cells (8–10). Consequently, the elevated AMP/ATP ratio activates adenosine monophosphate–activated protein kinase (AMPK) signaling pathway, which promotes cell cycle arrest and inhibits tumor cell proliferation (11). Recently, it has been demonstrated that clinical doses of metformin-bound PEN2 forms a complex with ATP6AP1, which leads to the inhibition of v-ATPase and the activation of AMPK without effects on cellular AMP levels (12). Preclinical studies and retrospective population-based studies have suggested antitumor activity of metformin alone or in combination with conventional anticancer drugs (13–16). Preclinical and clinical studies also have suggested that in patients with diabetes, prolonged use of metformin could be associated with a reduction of the HCC incidence (17, 18). Several research groups also have demonstrated metformin inhibited non-diabetes induced HCC in a mouse HCC model (19, 20). Williams *et al.* showed metformin prevents HCC development via alleviating p21 overexpression and ameliorating pro-tumorigenic microenvironment (19). Shankaraiah *et al.* showed metformin prevents hepatocarcinogenesis by attenuating fibrosis in a transgenic mouse model of HCC (20). However, despite decades of research, the mechanism by which metformin inhibits HCC is still highly debated. Additionally, the heterogeneity of HCC also impels us to determine the function and mechanism of metformin in chronic liver injury models of HCC.

Due to a deficiency in fumarylacetoacetate hydrolase (FAH) (21), hereditary tyrosinemia type 1 (HT1) is the severe inherited disorder of the tyrosine degradation. The FAH deficiency leads to the accumulation of toxic metabolites, mainly in the liver (22). Untreated HT1 patients usually die before the age of two. HT1 patients suffer multiple chronic complications, including cirrhosis and a high risk of HCC (22). NTBC/Nitisinone (a drug blocking the pathway upstream of FAH) greatly delay the morbidity of HT1 patients, but patients receiving NTBC may still developing HCC (23). The molecular basis of the pathogenic process in HT1 is still unclear. The murine model of *Fah*-deficiency (*Fah^-/-^*) is a suitable animal model, featuring all phenotypic and biochemical expressions of the HT1 patients (24). Moreover, the chronic liver injury model of *Fah^-/-^* mice closely resembles human alcohol-induced and c-myc-altered HCC (25). Therefore, the study of FAH deficient chronically injured mice can serve as a good indicator of human HCC.

In this study, we demonstrate that several unique signal pathways are vastly changed in an HCC model induced by chronic liver injury. These pathways are characterized by the regulation of glucose metabolic process and retinol metabolism. We find that the metformin increased the AMPK activity and subsequently reduced the HCC incidence. Our data also shows the metformin suppressed the *Cyp26a1* gene expression, leading to the increase of the all-trans-retinoicacid (atRA) level, and then reduced the number and ratio of pro-tumor CD8^+^ T cells. Moreover, metformin could suppress *Cyp26a1* gene expression through the AMPK/JNK/c-Jun pathway. Our results first disclose a link between the metformin and atRA, which may explain the commonness of their anti-tumor activities through inhibiting pro-tumor CD8^+^ T cells.

## Materials and methods

### Patient Specimens

Patient specimens were obtained from sample information service platform of Shanghai East Hospital under protocols approved by Shanghai East Hospital. Final diagnoses of the specimens were confirmed by experienced pathologists from Shanghai East Hospital. The study protocol conformed to the ethical guidelines approved by the Shanghai East Hospital Ethics Committee, and written informed consent was obtained from each patient.

### Analysis of public clinical datasets

The gene expression data of 7858 samples of 30 normal tissues (including liver organ) were downloaded from GTEx (https://gtexportal.org/home/), and the gene expression data of 710 samples of 17 para-tumor tissues and 7801 samples of 34 cancer types (including HCC) were downloaded from TCGA (https://portal.gdc.cancer.gov/). Correlation analyses between genes and signatures were calculated “cor()” function in R and also conducted in GEPIA database (http://gepia.cancer-pku.cn).

Moreover, we obtained mRNA expression from chronic injury liver cases from GSE89632 and GSE148355. To understand whether there is a similar trend that AMPK signaling pathway is inhibited in human chronic liver injury as in mouse model, we downloaded RNA-seq dataset GSE148355 (26) and Microarray dataset GSE89632 (27) from the public database GEO for verification. GSE148355 dataset was composed of HCC samples, “premalignant” samples with different degrees of fibrosis, and non-tumor samples from patients who had undergone surgical resection. We mainly used the normalized data (FPKM) of 62 premalignant and non-tumor samples for analysis, including 15 non-tumor samples (Normal), 10 low fibrosis samples (Fibrosis low, FL), 10 high fibrosis samples (Fibrosis high, FH), 10 cirrhosis samples (Cirrhosis, CS), 10 low degree of dysplasia nodule samples (Dysplastic nodule low, DL) and 7 high degree of dysplasia nodule samples (Dysplastic nodule high, DH). Likewise, we also analyzed microarray data from 63 NAFLD and healthy samples in GSE89632 dataset, including 20 simple steatosis samples (SS), 19 nonalcoholic steatohepatitis samples (NASH), and 24 healthy controls (HC).

In this study, we used the key genes in AMPK pathway to form a self-defined functional gene set, which was composed of PFKFB3, CPT1A, SIRT1, PPARGC1A, SLC2A4, PNPLA2, CRY1 and FOXO1. To infer whether AMPK pathway is inhibited during chronic liver injury, single-sample gene set enrichment analysis (ssGSEA) algorithm was used to calculate the enrichment score of this gene set in normal samples and samples with different degrees of chronic liver injury (28). Meanwhile, the expression of these eight single genes were also statistically analyzed in both healthy and chronic liver injury samples.

### Mouse model and Treatments

As described in previous studies (29, 30), 129S4 *Fah^−/−^* mice undergo liver failure and death. All mice were bred and maintained under specific-pathogen free conditions at Shanghai East Hospital. Mice were housed with a light: dark cycle of 12h, ambient temperature of 24℃ and humidity of 55%. Female mice were purely randomly allocated to metformin/atRA treated or untreated group. Mice in metformin experiments received metformin (B1970, APExBIO) via intraperitoneal injection at a dosage of approximately 250 mg/kg/day from 0 weeks until 12 weeks of chronic liver injury. 200 μg atRA (Sigma-Aldrich) dissolved in DMSO or DMSO alone was administered by intraperitoneal injection every other day from 0 weeks until 12 weeks of chronic liver injury. All experimental procedures on mice were approved by Institutional Animal Care and Use Committee of Shanghai East Hospital.

### Real time quantitative PCR (qRT-PCR)

Quantitative PCR was performed as described previously (29). Briefly, total RNA was isolated from liver of mice using RNAiso plus (TaKaRa). RNA was reverse-transcribed using the PrimeScript RT reagent Kit (TaKaRa). Transcript expressions were determined by qRT-PCR using SYBR Premix Ex Taq II (TaKaRa) and the QuantStudio^TM^ real-time PCR instrument (Applied Biosystems). The primers sequences used were as followed: Cyp26a1: 5’-AAGCTCTGGGACCTGTACTGT-3’ and 5’-CTCCGCTGAAGCACCATCT-3’, Gapdh: 5’-AGGTCGGTGTGAACGGATTTG-3’ and 5’-TGTAGACCATGTAGTTGAGGTCA-3’, Il6: 5’-GTCCTTCCTACCCCAATTTCC-3’ and 5’-TAACGCACTAGGTTTGCCGA-3’, Il1β: 5’-GCAACTGTTCCTGAACTCAACT-3’ and 5’-ATCTTTTGGGGTCCGTCAACT-3’, Tnf: 5’-CCCTCACACTCAGATCATCTTCT-3’ and 5’-GCTACGACGTGGGCTACAG-3’, Ltβ: 5’-TGGCAGGAGCTACTTCCCT-3’ and 5’-TCCAGTCTTTTCTGAGCCTGT-3’. qRT-PCR calculations were performed using the comparative CT method. The data are represented either as fold changes of the treated group compared to the control group or as 2^-*ΔΔ*CT^ mRNA transcript abundance.

### RNA-Seq and Transcriptomic Analyses of Mouse Liver Tissues

Total RNA (two liver tissues of C0 and C4, three liver tissues of C12 and Met12) were extracted using the mirVana miRNA Isolation Kit (Ambion) following the manufacturer’s protocol. RNA integrity was evaluated using the Agilent 2100 Bioanalyzer (Agilent Technologies, Santa Clara, CA, USA). The samples with RNA Integrity Number (RIN) ≥ 7 were subjected to the subsequent analysis. The libraries were constructed using TruSeq Stranded mRNA LTSample Prep Kit (Illumina, San Diego, CA, USA) according to the manufacturer’s instructions. Then these libraries were sequenced on the Illumina sequencing platform (HiSeqTM 2500 or Illumina HiSeq X Ten) and 125 bp/150 bp paired-end reads were generated.

The transcriptome sequencing and analysis were conducted by OE biotech Co., Ltd. (Shanghai, China). Raw data (raw reads) were processed using Trimmomatic (31). The reads containing ploy-N and the low-quality reads were removed to obtain the clean reads. Then the clean reads were mapped to reference genome using hisat2 (32). FPKM value of each gene was calculated using cufflinks, and the read counts of each gene were obtained by htseq-count (33). DEGs were identified using the DESeq (2012) R package functions estimate Size Factors and nbinomTest. “P value < 0.05” and “Fold Change > 2 or Fold Change < 0.5” was set as the threshold for significantly differential expression. Hierarchical cluster analysis of DEGs was performed to explore genes expression pattern. Gene Ontology (GO) enrichment and KEGG pathway enrichment analysis of DEGs were respectively performed using DAVID database(https://david.ncifcrf.gov/).

### Protein isolation and Western blot

Livers were harvested from normal *Fah*^-/-^ mice (100% NTBC), *Fah*^-/-^ mice with chronic liver injury (2.5% NTBC), or from different stages during the development of HCC. Lysates were prepared from the liver tissues using RIPA lysis buffer. The following primary antibodies were used for immunoblotting: anti-GAPDH (Protein tech, HRP-60004), anti-AMPK (CST, 2532), anti-P-AMPK (CST, 2535), anti-CYP26A1 (Santa Cruze, sc-53618), anti-P-JNK (CST, 9255), anti-JNK (CST, 9252), anti-P-c-Jun (Abcam, ab40476), anti-c-Jun (Abcam, ab32385). Primary and secondary HRP-labeled antibodies were used at 1:500 and 1:2000 dilutions respectively. Detection was performed with SuperSignal West Femto Maximum Sensitivity Substrate (Thermo Scientific).

### Metformin treatment of primary hepatocyte

Liver of chronic injury of *Fah*^-/-^ mice were perfused as before (34). 5x10^5^ cells of the primary hepatocyte were seeded in a 6-well plate (matrigel coated) with 2mL Advanced DMEM/F-12 (Thermo Fisher, 12634010) supplemented with 10% fetal calf serum. 24 hours later, the fresh medium replaces the old medium. The cells were treated with different concentrations of metformin or AICAR for 6 hours. In the presence of inhibitor, inhibitor was pre-treated to hepatocyte for 1 hour following the adding of metformin.

### Luciferase assay

Chronic injured liver of *Fah*^-/-^ mice were perfused. About 2x10^4^ cells of the primary hepatocyte were seeded in a 24-well plate with 1mL Advanced DMEM/F-12 supplemented with 10% fetal calf serum. 2 hours later, 1 mL Advanced DMEM/F-12 without fetal calf serum replaces the old medium. The second day, cells were transfected with a 6:1 ratio of a firefly luciferase reporter plasmid driven by a pGL3-RARE-responsive promoter (Addgene) and a Renilla luciferase reporter plasmid driven by a constitutive CMV promoter (Promega). 24 hours later, the culture medium was replaced with fresh Advanced DMEM/F-12 medium and metformin/atRA or vehicle. 24 hours later, activity of both reporters was measured using the Dual-Luciferase Reporter kit (Promega) and read on a Tecan Infinite 200 PRO Reader. The firely luciferase to renilla luciferase ratio is reported as “Firely/Renilla Luciferase Activity”.

### Serological Analyses

For Serum indicators, blood sample was collected from the retro-orbital sinus of test animals. Plasma was prepared using Microtainer plasma separator tubes (BD) and stored at −80 °C. Analysis the biochemical indicators of serum was according to the previously established protocol (29).

### Hematoxylin-Eosin staining (H&E), Immunohistochemistry (IHC)

Hematoxylin-Eosin staining (H&E) and Immunohistochemistry (IHC) are showed as before (29). For hematoxylin-eosin staining, fresh liver tissues were fixed with 4% paraformaldehyde (PFA), then were routinely embedded in paraffin and sectioned into slices (2 µm). The slices were processed for roasting, dewaxing and rehydration, and were stained with hematoxylin (Beyotime, Shanghai, China) for 5– 10 min, rinsed with water for 15 min, sliced into 95% alcohol (Sinoreagent, Shanghai, China) for 30 seconds, and then stained with eosin (Beyotime, Shanghai, China) for an appropriate amount of time (0.5–2 min). Finally, the slice is dehydrated rapidly and mounted with neutral resin (MXB, Fujian, China).

For immunohistochemistry staining, IHC steps are the same as H&E before rehydration. The slices were soaked in 0.01 M citric acid buffer (pH 6.0) or EDTA (pH 9.0) and placed in a pressure cooker for 2–4 min at 121 °C/100 kpa. Cool to room temperature. 3% H2O2solution blocked endogenous peroxidase and 1% BSA blocked nonspecific loci for 30 min at room temperature. The slices were incubated with primary antibodies for at 4 °C overnight and secondary antibodies conjugated with HRP at 37 °C for 30 min. Staining with DAB (Vector Laboratories, Burlingame, CA) was applied to the sections. The sections were stained with hematoxylin (Beyotime, Shanghai, CHN), dehydrated rapidly and mounted with neutral resin (MXB, Fujian, CHN). The used primary antibodies are summarized as below: CD8 (Abcam, ab209775), CD45 (CST, 70257).

### Flow Cytometry

Fresh liver tissue samples were minced, then digested in mouse Liver Dissociation Kit (MACS 130-105-807) for 0.5h at 37°C. Cell suspensions were generated through a 70μm nylon mesh. After filtration and washing, cells were suspended in a 40% Percoll solution following by centrifugation at 600 g for 20 min. Cells were prepared as a single-cell suspension for FACS staining. For surface staining, the following antibodies were used: APC-CY7-LIVE/DEAD, FITC-CD45, V450-CD3, AF700-CD4, V500-CD8, BV605-CD49B, PE-F4/80, BB700-CD11B. The stained cells were acquired for analysis using the Cytoflex (BECHMAN). Flow cytometry data was analyzed with FlowJo software (Tree Star Inc.).

### Blood Glucose Measurement

Blood was sampled in mice by nicking the tail vein and blood glucose levels were measured using ACCU-CHEK Active test strips read by an ACCU-CHEK Active meter (Roche Diagnostics, Indianapolis, IN) following the manufacturer’s instructions.

### Single cell RNA sequencing (scRNA-seq) analysis

The scRNA-seq data of *Fah^-/-^* mouse at 0, 12 and 18 weeks after the withdrawal of NTBC were downloaded from GEO databases with the GEO accession number GSE130880 (35). The dimension reduction analysis of Stochastic Neighbor Embedding (t-SNE) and Uniform Manifold Approximation and Projection (UMAP) were calculated by “RunTSNE()” and “RunUMAP()” function of Seurat package (R-version 4.0.5). Correlation analyses between signatures were calculated “cor()” function in R. In addition, we used ssGSEA algorithm (28) to calculate the scores of pro-tumor CD8^+^ T cell signature in CLI *Fah^-/-^* mouse at 0, 12 and 18 weeks.

### Statistics

All results were presented as “mean ± SEM” as indicated. The differences between groups were analyzed with one-way or two-way ANOVA, Student’s t-test or Mann-Whitney U test with two-tailed. The overall survival (OS) and disease-free survival (DFS) were analysis by Log-Rank test using GraphPad Prism 7. ** P < 0.01, * P < 0.05, NS = not significant.

### Data Availability

The RNA sequencing raw data for this study were generated at OE biotech Co., Ltd. All data generated or analysed during this study are included in the manuscript and supporting file; Source Data files have been provided for Figures 1 and 3.

**Figure 1.**
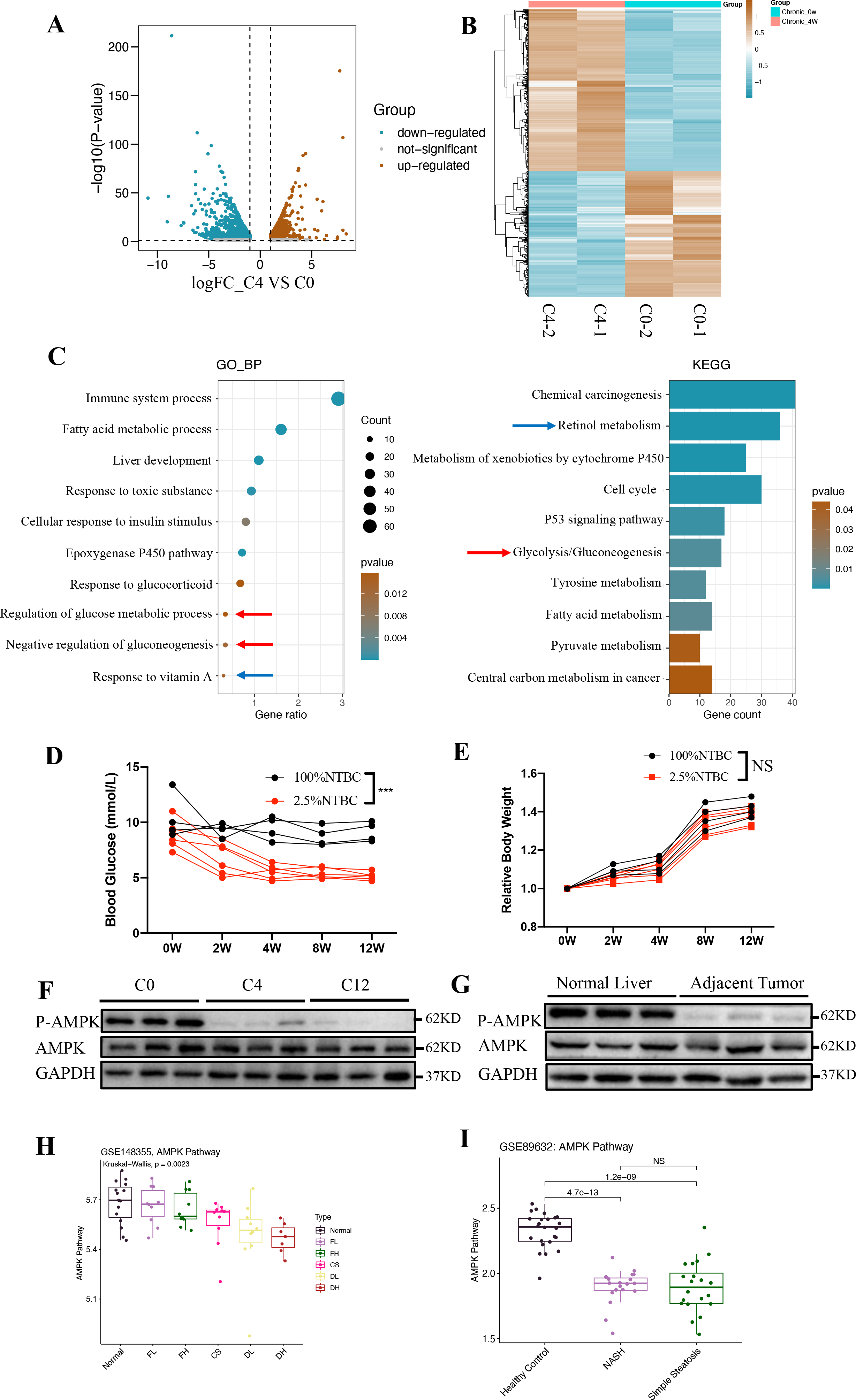
Characteristics of *Fah^-/-^* mice with chronic liver injury. (A) RNA sequencing showing volcano plot of differential expression analysis results from 4 weeks of chronic liver injury (C4) vs normal *Fah^-/-^* mouse liver (C0). (B) Heat map illustrates differential gene expression of liver tissue from 4 weeks chronic liver injury (C4, labeled as FT4 here) vs normal *Fah^-/-^* mouse (C0, labeled as FT0 here). (C) Gene Ontology (GO) classification, and Kyoto Encyclopedia of Genes and Genomes (KEGG) pathway enrichment analyses of the differentially expressed genes from 4 weeks chronic liver injury (C4) vs normal *Fah^-/-^* mouse liver (C0). (D) Body weight (BW) in *Fah^-/-^* mice with chronic liver injury (2W, 4W, 8W and 12W) and normal *Fah^-/-^* mouse (0W). (E) Blood glucose levels in *Fah^-/-^* mice with chronic liver injury (2W, 4W, 8W and 12W) and normal *Fah^-/-^* mouse (0W). (F) Western blot analysis of AMPK activity from 4 weeks (C4), 12 weeks (C12) chronic liver injury and normal *Fah^-/-^* mouse liver (C0). (G) Western blot analysis of P-AMPK level in healthy human liver tissues and adjacent noncancerous liver tissues of HCC patients. (H) Enrichment score of AMPK pathway in normal and premalignant tissues. The enrichment score of AMPK pathway was calculated by ssGSEA algorithm. (I) Enrichment score of AMPK pathway in normal tissues and NAFLD tissues. Normal: non-tumor Normal control, FL: low Fibrosis, FH: high Fibrosis, CS: Cirrhosis, DL: Dysplastic nodule Low, DH: Dysplastic Nodule high, NASH: non-alcoholic steatohepatitis. All data represented as “mean ± SEM”. Statistical significance was determined by unpaired two-tailed t-test. *P < 0.05; **P < 0.01; ***P < 0.001.

## Results

### Characters of *Fah^-/-^* mice with chronic Liver injury

Using the *Fah^-/-^* mouse model as a surrogate for chronic liver disease, we first described the liver phenotype of *Fah^-/-^* mice under different conditions with our and other published works (29, 36). As detailedly described in Figure S1, under 100% concentration of NTBC (7.5 mg/L), the liver of *Fah^-/^*^-^ mice was preserved normally; *Fah^-/-^* mice developed acute liver injury and died at 3-6 weeks without any NTBC; *Fah^-/-^* mice survived under 2.5% concentration of NTBC (0.2 mg/L), but suffered chronic liver injury and formed HCC after 12 weeks.

To gain further insight into the molecular mechanisms that contribute to HCC development in chronic liver injury, we first aimed to characterize the dynamic transcriptomic changes in chronic injured liver compared with normal liver. After performing high throughput RNA Sequencing, the differentially-expressed genes between normal livers of *Fah^-/-^* mice (100% NTBC, named as C0) and chronic injured livers of *Fah^-/-^* mice (2.5% NTBC for 4 weeks, named as C4) were identified (Fig. 1A-B, Table S1). Two thousand genes with significantly altered expression were identified (log2 Fold Change > 1, P value < 0.05). Genes with expression level altered by more than two-fold were further analyzed by gene enrichment analyses, including Gene Ontology (GO) and KEGG. Several pathways had been enriched in *Fah^-/-^* mice with Chronic Liver Injury (CLI *Fah^-/-^* mice) (Fig. 1C, Table S2). HCC represents a classic instance of inflammation-related cancer, and chemically or genetically induced HCC is highly dependent on inflammatory signaling (3). We found many of these differentially expressed genes are involved in inflammatory processes, and they are also implicated in HCC development (Fig. 1C). Consistently, several KEGG pathway terms, especially the p53 signaling pathway, were altered in *Fah^-/-^* mice livers (Fig. 1C). The process of retinol metabolism was also significantly altered both in the CLI and normal *Fah^-/-^* mice (Fig. 1C, blue arrow). Of note, the glucose metabolic process was also enriched in the CLI *Fah^-/-^* mice according to GO and KEGG analyses (Fig. 1C, red arrow). Body weight and blood glucose level in the CLI *Fah^-/-^* mice were measured. We found there is similar body weight and significantly lower blood glucose level in the CLI *Fah^-/-^* mice compared with normal *Fah^-/-^* mice, indicating that HCC model on the CLI *Fah^-/-^* mice represents non-diabetes induced HCC (Fig. 1D-E). The function of metformin depends on the activation of AMPK and thereby we wanted to know whether the AMPK activity was changed in our chronic liver injury model of *Fah^-/-^* mice. We detected the activity of AMPK in *Fah^-/-^* mice livers of C0 (Uninjured Control, 0 week), C4 (CLI, 4 weeks) and C12 (CLI, 12 weeks), respectively. And we found that the activity of AMPK in liver decreased significantly when the *Fah^-/-^* mice suffered CLI (Fig. 1F). We tested human patients with normal or cirrhotic liver tissues and found that the AMPK activity is also repressed in chronic injured human liver (Fig. 1G). To understand whether there is a similar trend on the inhibition of AMPK signaling between the human chronic liver injury and the mouse model, we obtained two human chronic liver injury related datasets from GEO database, and ssGSEA algorithm was applied to evaluate whether the AMPK pathway was hampered under chronic liver injury. It was found that with the aggravation of liver fibrosis, the enrichment score of AMPK pathway gradually declined, and the AMPK pathway tended to be suppressed in all status of chronic injured liver compared with normal tissues (Fig. 1H). Similarly, the enrichment score of AMPK pathway in Non-Alcoholic Steatosis Hepatitis (NASH) and simple steatosis groups were significantly lower than those in the healthy control group (Fig. 1I). In addition, at the single gene level, we observed that the expression level of FOXO1, CRY1 and other AMPK pathway core genes were significantly lower in chronic injured liver than those in healthy control (Fig. S2). Beside of these data, preclinical and clinical studies have suggested the activity of AMPK in human liver decreased significantly with chronic liver injury and lower activity of AMPK indicates poorer survival (37–40). All the data above showed that the CLI *Fah^-/-^* mice have signal pathway altering patterns mimicking human chronic liver diseases, and is a non-diabetic induced HCC mouse model serving as a good indicator of human HCC.

### Metformin prevents hepatocarcinogenesis in *Fah^-/-^* mice with chronic liver injury

Previous studies suggest that metformin treatment inhibits HCC development in several animal models of HCC (19, 20). We tried to figure out whether metformin inhibited HCC formation of the CLI *Fah^-/-^* mice. To investigate whether the metformin affects HCC formation of the CLI *Fah^-/-^* mice, we switched from 100% to 2.5% NTBC administration at the experimental duration of 8 weeks, at the same time, animals were treated with daily intraperitoneal injections of either metformin (Met12, 250 mg/kg body weight) or an equal volume of saline solution (C12) for 12 weeks (Fig. 2A). The main pharmacologic mechanism of metformin is activating the AMPK pathway (11). The results of western blot showed that in our chronic liver injury model, AMPK was activated in the metformin treatment group, indicating that the metformin worked (Fig. 2B). HCC were obviously visible in both macroscopic and histological examination in all *Fah^-/-^* mice (n=12). Metformin treatment significantly delayed tumor formation; only 50% of the *Fah^-/-^* mice developed HCC after 12-week 2.5% NTBC treatment (n=12) (Fig. 2C). Furthermore, *Fah^-/-^* livers without metformin treatment displayed a significantly greater number and size of tumors than those treated with metformin (Figs. 2D-E). In addition, analyses on serum liver function indicators were used to confirm the degree of liver injury in both groups. We found the level of serum aspartate transaminase (AST), serum alanine transaminase (ALT), and the AST/ALT ratio were accordingly not significantly changed in both groups (Fig. 2F). Taken together, these data indicate that metformin prevents hepatocarcinogenesis in *Fah^-/-^* mice with chronic liver injury, even without alleviating liver functions.

**Figure 2.**
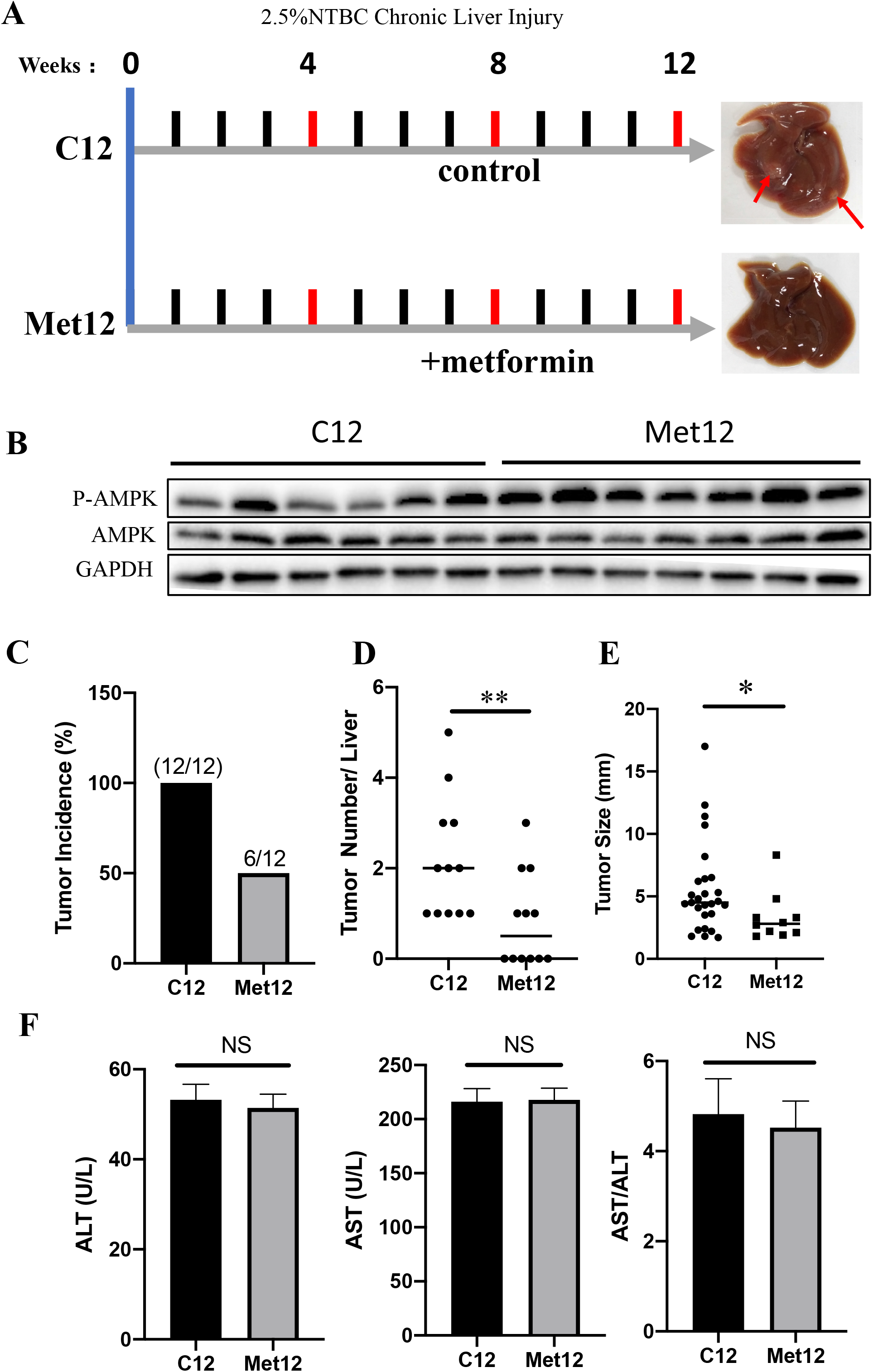
Effects of metformin on the characteristics of precancerous livers and HCC incidence in *Fah^-/-^* mice. (A) Schematic diagram showing the experimental set. Representative photographs of livers about chronic liver injury for 12 weeks without metformin (C12) and with metformin (Met12) are showing. (B) Western blot analysis of P-AMPK and AMPK protein level in liver tissues of C12 and Met12. (C) Graphs representing tumor incidence of *Fah^-/-^* mice with and without metformin (Met12 and C12). (D-E) Scatter plots displaying the tumor numbers (D) and size of tumors (E) in Fah-deficient livers at C12 and with Met12. (F) Serological indexes of liver injury (ALT, AST and AST/ALT) are almost not changed in the serum of *Fah^-/-^* mice at C12 and Met12. *P < 0.05; **P < 0.01; ***P < 0.001; NS: not significant.

### Metformin suppressed *Cyp26a1* expression of hepatocyte *in vivo* and *in vitro*

The major function of metformin depends on the activation of the AMPK pathway (11). To identify potential AMPK-targeted gene that mediating the metformin-induced hindering of HCC development in *Fah^-/-^* mice, RNA-Seq and corresponding differential gene expression analyses on the livers of *Fah^-/-^* mice, with or without metformin treatment, was performed. The results revealed that the mRNA levels of 200 genes were significantly changed with metformin treatment (log2 Fold Change >1) (Fig. 3A, Table S3). From these 200 altered genes, we presumed the gene *Cyp26a1* was the target of metformin in the hindering of hepatocarcinogenesis for four reasons: (i) Williams *et al.* showed metformin prevents HCC development in the chronic liver injury of *Ncoa5*^+/−^ mice (19). We comparatively re-analyzed our and Williams’s sequencing data (GSE110524). There are 200 significantly changed genes in our RNA-Seq data (127 upregulated genes, 73 downregulated genes), 157 significantly differential genes in GSE110524 data (53 upregulated genes, 104 downregulated genes) (19). We conducted intersection analyses of the up-regulated and down-regulated genes between these two data, respectively. There was no up-regulated overlapped genes. Interestingly, only one intersected gene, *Cyp26a1*, was obtained in the down-regulated genes by metformin (Fig. 3B). (ii) The expression of *Cyp26a1* is significantly up-regulated in C4 compared to C0, and the decrease of its expression significantly affects the physiological and biochemical function of liver (Table S1). (iii) *Cyp26a1* encodes one of the cytochrome P450 superfamily enzymes which metabolize all-*trans*-retinoic acid (atRA), 9-*cis*RA and 13-*cis*RA (41). The decrease of *Cyp26a1* leads to the augmentation of atRA level, while *at*RA has been demonstrated as anti-tumor metabolite in several tumor types (42–45). (iv) In the process of chronic liver injury of *Fah^-/-^* mice, the metabolic signal pathway of retinol (a precursor of atRA) was enriched by GO and KEGG analysis (Fig. 1C blue arrow), which implies retinol pathway is important in the HCC formation. For all these reasons, we presume metformin may suppressed the expression of *Cyp26a1* to prevent hepatocarcinogenesis in *Fah^-/-^* mice with chronic liver injury.

**Figure 3.**
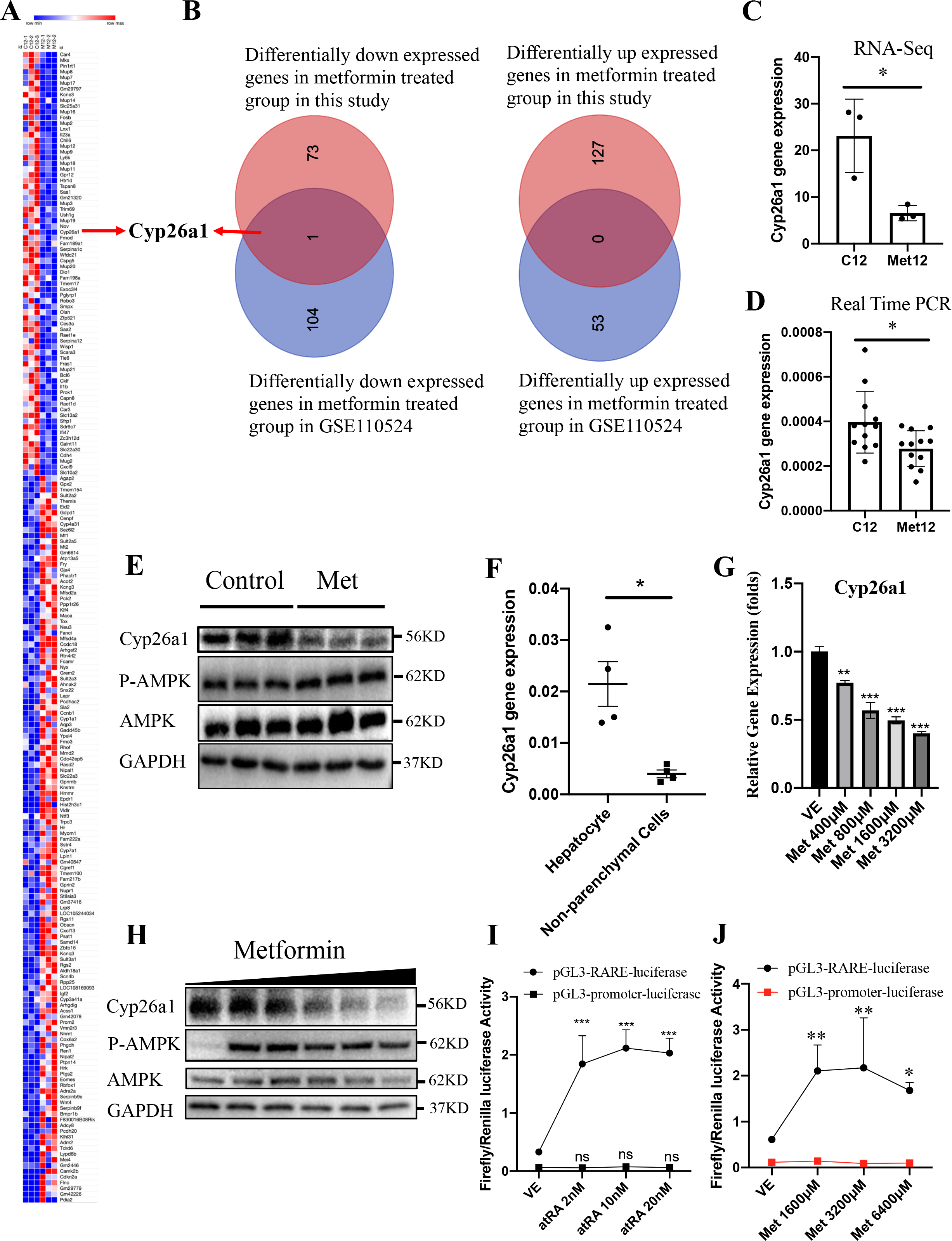
Effects of metformin on *Cyp26a1* Gene in mouse hepatocyte. (A) Differentially up- or down-regulated genes between C12 and Met12. (Log2 Fold Change >1, FDR<0.05). (B) The intersection of our sequencing results and GSE110524. (C) RNA-seq data of *Cyp26a1* gene. (D) Quantitative analysis of *Cyp26a1* gene in liver tissues from C12 and Met12 was performed by qRT-PCR. (E) Western blot analysis of relative protein in liver tissues from C12 and Met12. (F) qRT-PCR analysis of *Cyp26a1* gene expression between mouse hepatocyte and non-parenchymal cells. perfused hepatocyte and non-parenchymal cells were separated by low-speed centrifugation. (G) Perfused hepatocyte treated with metformin in vitro. qRT-PCR analysis results of *Cyp26a1* gene expression with different concentrations of metformin. (H) Western blot analysis of *Cyp26a1* gene expression in perfused adherent hepatocyte with different concentrations of metformin *in vitro*. (I) RARE luciferase assay on *Fah^-/-^* hepatocyte cultured with increasing amounts of atRA. Data representative of 3 independent experiments. Two-way ANOVA was used. (J) RARE luciferase assay on *Fah^-/-^* hepatocyte cultured with increasing amounts of metformin. Data representative of 3 independent experiments. Two-way ANOVA was used. Data are represented as “mean ± SEM”. *P < 0.05; **P < 0.01; ***P < 0.001; NS: not significant.

CYP26A1 functions primarily to eliminate bioactive atRA (46). RNA-seq data showed *Cyp26a1* gene expression in metformin treatment group was reduced (Fig. 3C). In line with the results from the RNA-Seq analysis, *Cyp26a1* mRNA levels were down-regulated in metformin-treated liver tissues of *Fah^-/-^* mice according to the qRT-PCR assay (Fig. 3D). Western blot also showed CYP26A1 protein expression was reduced in metformin-treated liver tissues (Fig. 3E). In the adult, *Cyp26a1* is mostly highly expressed in the liver (47). We examined the cellular source of *Cyp26a1* in the mouse liver. Perfused mouse liver was separated to hepatocytes and non-parenchymal cells by low-speed centrifugation (50 g for 2 min). qRT-PCR revealed that *Cyp26a1* was mainly expressed by hepatocyte in the mouse liver (Fig. 3F). The scRNA-seq data of immune cells of another HBV-induced HCC model in Fah^-/-^ mouse (35) also showed that only 23 of 27058 immune cells expressed *Cyp26a1* (data not shown). Next, we examined whether metformin directly suppressed the *Cyp26a1* expression in hepatocytes. Perfused hepatocytes were plated on a plate containing matrigel. qRT-PCR and western blot showed that the expression of *Cyp26a1* in hepatocytes decreased with the increase of metformin concentration (Fig. 3G-H). Protein CYP26A1 plays a major role in atRA clearance which means down-regulation of *Cyp26a1* leads to the elevation of atRA level (41). The pGL3-RARE-luciferase is a plasmid contains retinoic acid receptor (RARE) response element in front of luciferase (firefly) reporter gene, which can reflect the level of atRA (45, 48). We transfected this plasmid and control plasmid (pRL-CMV-renilla-luciferase) to primary hepatocytes of the *Fah^-/-^* mice with chronic liver injury. The Firely/renilla luciferase activity of pGL3-RARE-luciferase is increased with atRA treatment while pGL3-promoter-luciferase is unchanged (Fig. 3I), which means pGL3-RARE-luciferase activity could reflect the level of atRA in hepatocytes. We then tested pGL3-RARE-luciferase and pGL3-promoter-luciferase activity on the metformin treatment. We found that the Firely/renilla luciferase activity of pGL3-RARE-luciferase is increased while pGL3-promoter-luciferase is unchanged with metformin treatment, which means metformin increased the level of atRA (Fig. 3J). Together, these data demonstrated metformin directly suppresses the *Cyp26a1* gene expression of hepatocytes and thereby leads to the elevated level of atRA, thereafter exerting anti-tumor effects.

### Metformin decreased the expression of *Cyp26a1* through the AMPK/JNK/c-Jun signaling

The AMPK is a major effector of metformin (49), and acadesine (AICAR) is another AMPK activator (50). We treated perfused primary adherent hepatocytes of *Fah^-/-^* mice with different concentration of AICAR. qRT-PCR and western blot showed the expression of *Cyp26a1* was also suppressed by AICAR (Fig. 4A-B). We speculated that the AMPK may be involved i n the decreased expression of *Cyp26a1* induced by metformin and AICAR. Compound C (dorsomorphin) is an effective, reversible and selective AMPK inhibitor, and is widely used to imply AMPK-dependence experimentally (49). Compound C (80 μM) attenuates the ability of metformin in activating AMPK in mouse hepatocytes (Fig. 4D). qRT-PCR and western blot showed Compound C attenuates the ability of metformin in suppressing the *Cyp26a1* expression which means the AMPK is a major effector of metformin in suppressing *Cyp26a1* gene expression (Fig. 4C-D).

**Figure 4.**
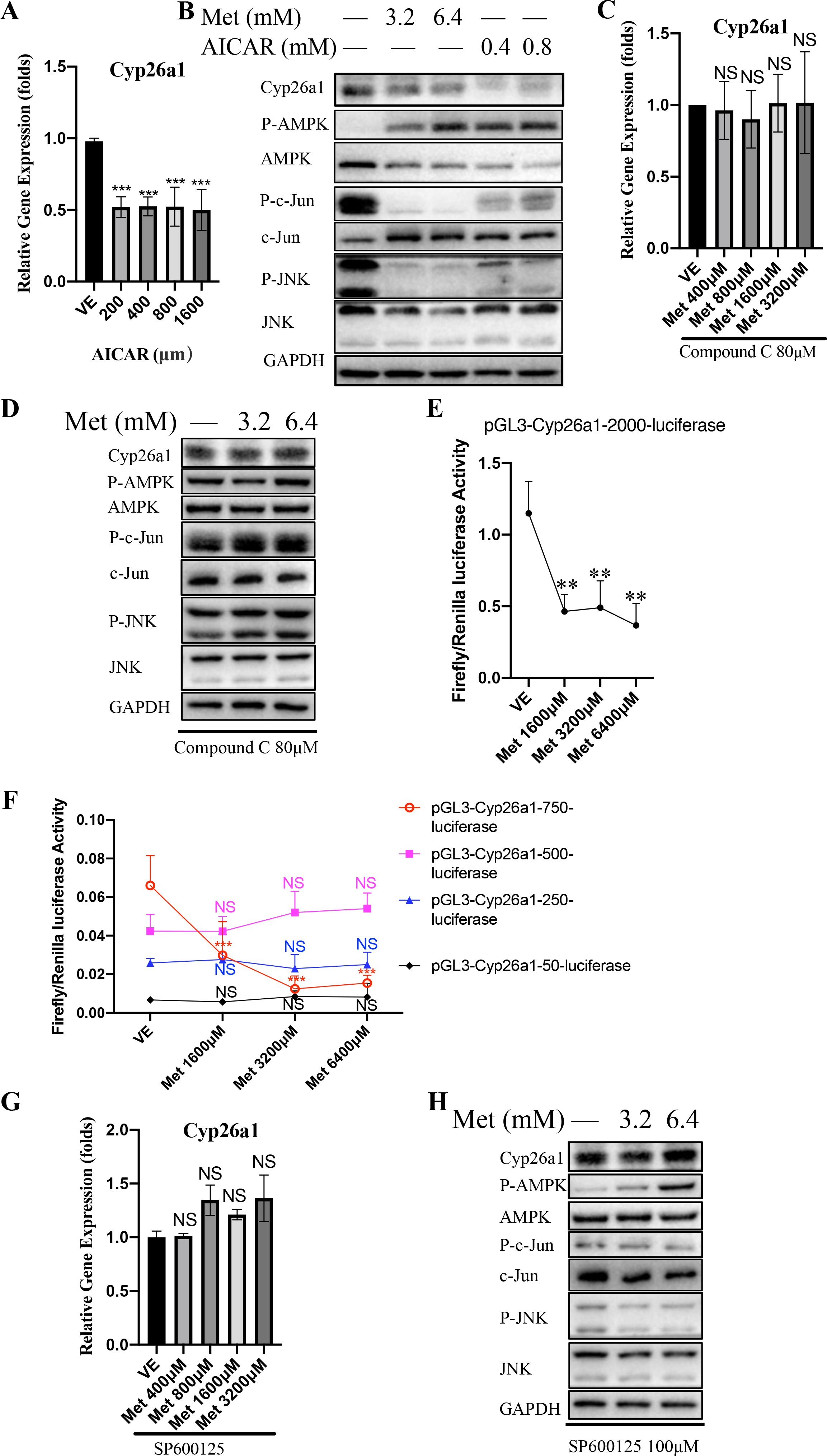
Metformin suppressed *Cyp26a1* gene by inhibiting JNK/c-Jun pathway. (A) qRT-PCR analysis results of *Cyp26a1* gene expression of primary hepatocytes with different concentrations of AICAR. (B) Western blot analysis of relative gene expression in perfused adherent hepatocyte with different concentrations of metformin and AICAR *in vitro*. (C) qRT-PCR analysis results of *Cyp26a1* gene expression. Primary hepatocytes treated with AMPK inhibitor Compound C (80 μM) in the absence or presence of metformin for 6 hours. Treatment with the inhibitor started 1 h before metformin treatment. (D) Western blot analysis of relative gene expression. Primary hepatocytes treated with AMPK inhibitor in the absence or presence of metformin for 6 hours. Treatment with the inhibitor started 1 h before metformin treatment. (E) We transfected plasmid (pGL3-cyp26a1-promoter-2000bp-luciferase) into primary hepatocyte cultured with increasing amounts of metformin. Data representative of 3 independent experiments. T wo-way ANOVA was used. (F) The truncated mouse *Cyp26a1* promoter mutation plasmids were transfected into primary hepatocyte cultured with increasing amounts of metformin, separately. Data representative of 3 independent experiments. Two-way ANOVA was used. (G) qRT PCR analysis results of *Cyp26a1* gene expression. Primary hepatocytes treated with JNK inhibitor SP 600125 (100 μM) with increasing amounts of metformin for 6 hours. Treatment with the inhibitor started 1 h before metformin treatment. (H) Western blot analysis of relative gene expression. Primary hepatocytes treated with JNK inhibitor SP600125 (100 μM) with increasing amounts of metformin for 6 hours. Treatment with the inhibitors started 1 h before metformin treatment. *P < 0.05; **P < 0.01; ***P < 0.001; NS: not significant.

To determine whether metformin mediated *Cyp26a1* expression through the repression of its promoter, we constructed plasmid pGL3-cyp26a1-promoter-2000bp-luciferase, which was generated by using the promoter of mouse *Cyp26a1* gene (- 2000 to +60 relation to transcription start site) to replace SV40 promoter of pGL3-promoter-luciferase. The mouse *Cyp26a1* promoter activity was significantly suppressed after metformin treatment (Figure 4E) while pGL3-promoter-luciferase is not changed (Fig. 3J), which means metformin-suppressed *Cyp26a1* expression is mediated through the repression of its promoter.

To define the roles of the *cis*-regulatory elements of the *Cyp26a1* promoter in response to metformin regulation, a series of truncated mutants of the *Cyp26a1* promoter were generated (-750 to +60, -500 to +60, -250 to +60 and -50 to +60). pGL3-cyp26a1-promoter-750bp-luciferase also showed significantly decreased luciferase activity by metformin. The shorter truncated mutants (-500 to +60, -250 to +60 and -50 to +60) of the *Cyp26a1* promoter blocked metformin-decreased *Cyp26a1* promoter activity, indicating that the sequence between nucleotide -750 and -500 was critical for the suppression of the *Cyp26a1* by metformin treatment (Fig. 4F). We predicted the transcription factors of this sequence (nucleotide -750 to -500 relation to transcription start site) using PROMO database (http://alggen.lsi.upc.es/cgibin/promo_v3/promo/promoinit.cgi?dirDB=TF_8.3). There are forty predicted transcription factors (Fig. S3), and among them a predicted c-Jun binding site is located in this region. Previous study showed the JNK/c-Jun signal pathway could be regulated by metformin-AMPK pathway (51). We found P-JNK and P-c-Jun was inhibited by metformin in hepatocytes (Fig. 4B). When activity of AMPK was pharmacologically inhibited by Compound C, such reduction by metformin treatment in *Cyp26a1*, P-JNK and P-c-Jun expression were not observed (Fig. 4D). Furthermore, we used JNK inhibitor SP600125 to determine whether metformin-decreased *Cyp26a1* expression is mediated through the repressed of JNK. In the presence of SP600125, JNK activity is suppressed and metformin lost the effect of inhibiting JNK because JNK activity has been inhibited in advance by SP600125. In parallel, *Cyp26a1* could not be decreased by metformin even AMPK is activated when JNK activity was not suppressed by metformin (Figs. 4G-H). Taken together, the results demonstrate that the *Cyp26a1* expression is inhibited by metformin through the AMPK/JNK/c-Jun signal pathway.

### atRA supplementation decreases HCC formation in chronic liver injury model

Metformin could promote the level of atRA through downregulation of *Cyp26a1* gene expression (Fig. 3). The atRA is usually considered as an anti-tumor small molecule (41). We wanted to study the antitumor effect of atRA in our chronic liver injury model. IP injection of atRA was administered to *Fah^-/-^* mice at the beginning of chronic liver injury (Fig. 5A). Tumor incidence, tumor number and tumor diameter in atRA-treated mice were significantly reduced compared to vehicle-treated mice, respectively (Fig. 5B-D). We also found that the levels of serum aspartate transaminase (AST), serum alanine transaminase (ALT), and AST/ALT ratio were accordingly not significantly changed in atRA treatment compared to untreated control (Fig. 5E). These data demonstrate a robust therapeutic benefit conferred by atRA supplementation in our chronic liver injury model.

**Figure 5.**
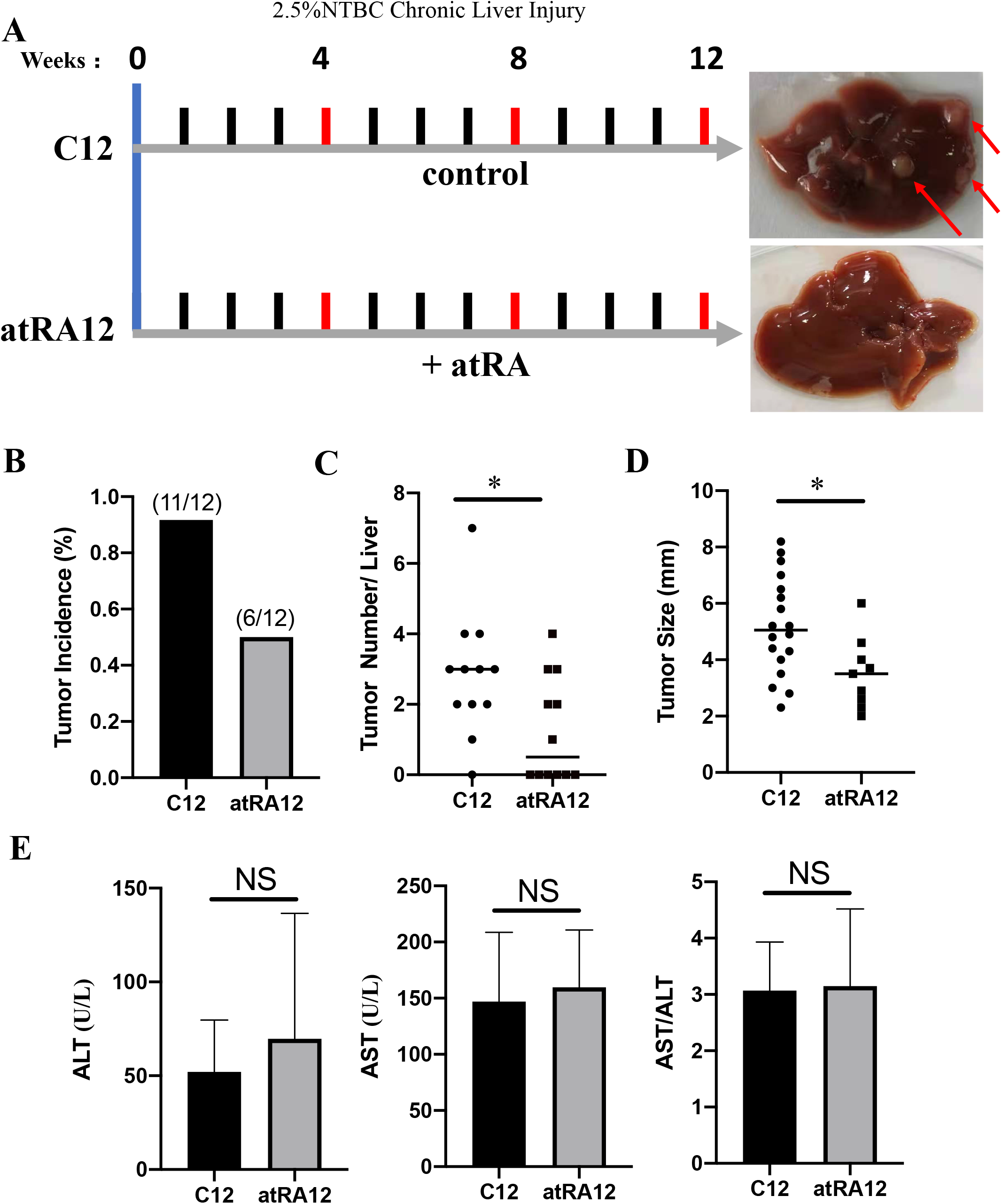
Effects of atRA on HCC formation in *Fah^-/-^* mice with chronic liver injury. (A) Schematic diagram showing the experimental set. Representative photographs of livers about chronic liver injury for 12 weeks without atRA (C12) and with atRA treatment (atRA12) are showing. (B) Graphs representing tumor incidence of *Fah^-/-^* mice with and without atRA treatment. (C) Scatter plots displaying the tumor numbers of *Fah^-/-^* mice with and without atRA treatment. (D) Scatter plots displaying the tumor size in *Fah^-/-^* mice between C12 group and atRA12 group. (E) Indicators of hepatocyte injury (ALT, AST and AST/ALT) are almost not changed in the serum of *Fah^-/-^* mice at C12 and atRA12 groups. *P < 0.05; **P < 0.01; ***P < 0.001; NS: not significant.

### Metformin and atRA both reduce CD8*^+^* T cell number and ratio in CLI mouse model

Recent publications have shown that metformin is able to modulate the interaction between tumor cells and their microenvironment and thus presenting an immune-mediated antitumor effect (52–55). In addition, the antitumor properties of atRA are also closely related to immune cells (45). To explore the mechanism behind tumor suppression, we studied immune cell subsets in the CLI *Fah^-/-^* mice on metformin or atRA treatment. A prominent expansion of the proportion of hepatic CD8^+^ cells among the hepatic CD45^+^ cells was found in C12 mice when compared with C0 mice, and the proportion of CD8^+^ T cells were significantly reduced in metformin-treated mice (Met12) compared with C12 mice, whereas no changes were found in other immune subsets (Kupffer cells, CD4^+^ T, NK and CD3^+^ cells) (Fig. 6A-B). Although the proportion of CD4^+^ in all CD45^+^ cells were significantly increased in metformin-treated mice (Met12) compared with C12 mice, there is no change between C12 and C0 mice, so we focused on the role of CD8^+^ T cells in the metformin-induced tumor formation hindering. A similarity result was found by IHC staining, chronic injured liver (C12) contained much more CD45^+^ cells and CD8^+^ T cells than the liver of mice on 100% NTBC (C12 VS C0), the number of CD8^+^ T cells were reduced by metformin treatment while the number of CD45^+^ cells remained unchanged (Met12 VS C12) (Figs. 6C-D). Accordingly, the number of hepatocytes per field significantly dropped in the histological sections of the CLI *Fah^-/-^* mice compared with normal liver tissue sections, which hinted that the size of hepatocytes enlarged significantly in the CLI *Fah^-/-^* mice compared with those in normal liver (C12 VS C0) (Fig. 6D). Hepatic CD8^+^ T cells are integral to antitumor immunity via direct antigen-specific cytotoxic targeting of tumors. However, recent researches showed that the CD8^+^ T cells contribute to HCC tumor formation in chronic liver injury model (19, 25, 56, 57). These papers showed the CD8^+^ T cells secrete pro-tumor cytokines (IL-1β, IL-6, Tnfa and Ltβ) in the process of chronic liver injury. Especially, CD8^+^ T cells as pro-tumor immune cell has been demonstrated in CLI *Fah^-/-^* mice, although the adopted mouse model is a little different to our CLI *Fah^-/-^* mice (25). We aimed to delineate which cytokine mediates the impact of CD8^+^ T cells on hepatocarcinogenesis in our disease model. P ro-tumor cytokines mRNA levels were measured by qRT-PCR, and the induction of Tnfa and Ltβ strikingly correlated with tumor development in the CLI *Fah^-/-^* mice. Moreover, chemokines like the Tnfa and Ltβ were significantly reduced in metformin treatment (Fig. 6E). Together, our results indicate that metformin could reduce the number and ratio of CD8^+^ T cells to suppress HCC formation in the CLI *Fah^-/-^* mice.

**Figure 6.**
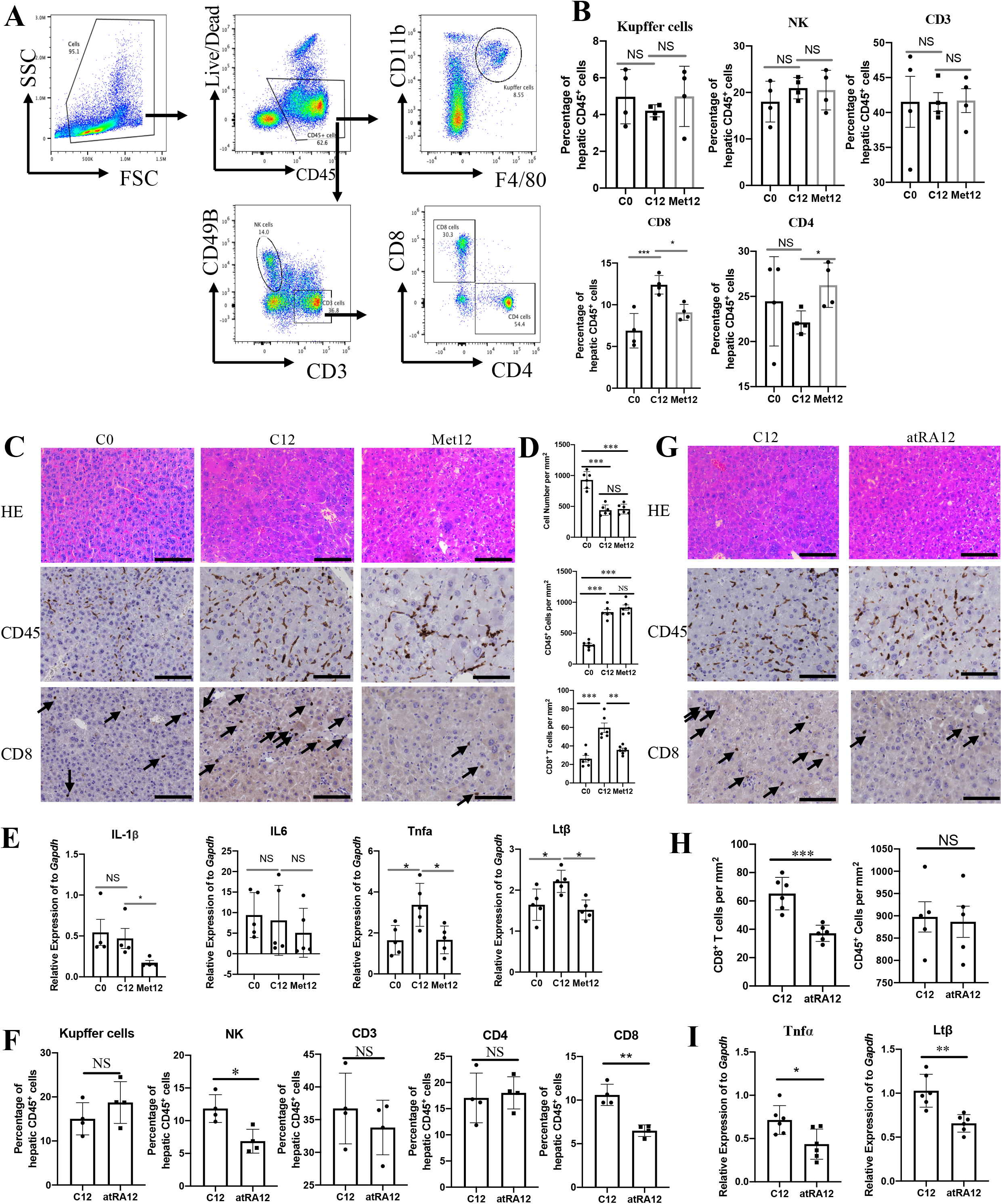
Effects of Metformin and *at*RA on the characteristics of immune cells in *Fah^-/-^*mice. (A) Single immune cells were stained with anti-CD3, CD8, CD45, CD4, CD49B, F4/80, CD11B and Live/Dead antibodies. The phenotype of liver immune cells was analyzed by FACS. (B) Liver infiltrating immune cells were measured treated with or without metformin. Data represent “mean ± SEM” of four pooled experiments. Statistical significance was determined by unpaired two-tail t-test. (C) Representative pictures of the indicated H&E staining and immunohistochemistry staining. Scale bar, 50μm. (D) Quantification of hepatocyte cell number, CD45^+^ cell number and CD8^+^ cell number per square millimeter. (E) Quantitative analysis of pro-tumor cytokines of liver tissues. (F) Liver infiltrating immune cells were measured treated with or without atRA. Data represent “mean ± SEM” of four pooled experiments. Statistical significance was determined by unpaired two-tail t-test. (G) Representative pictures of the indicated H&E staining and immunohistochemistry staining. Scale bar, 50μm. (H) Quantification of CD45^+^ cell number and CD8^+^ cell number per square millimeter in the liver. (I) qRT-PCR analysis of pro-tumor cytokines of liver tissues. *P < 0.05; **P < 0.01; ***P < 0.001; NS: not significant.

Next, we sought to clarify the mechanism that was responsible for the anti-tumor effect of atRA. Since CD8^+^ T cells play a crucial role in pro-tumor immunity in our model, we examined the CD8^+^ T cell and other immune cells in the atRA-treated CLI *Fah^-/-^* mice. A significant decrease in the percentage of CD8^+^ T cells was found after atRA treatment compared with vehicle treatment (atRA12 vs C12) (Fig. 6F). Besides, a significant decrease in the percentage of NK cells was found after atRA treatment while was not found after metformin treatment (Fig. 6B). There were no changes in other immune subsets after atRA treatment compared with vehicle treatment (Kupffer cells, CD4^+^ T and CD3^+^ cells). Different effects between metformin and atRA treatment indicate that the signal pathways regulated by metformin and atRA are not completely consistent in our mouse model. IHC staining also showed that with atRA treatment, the number of CD8^+^ T cells decreased significantly while the number of CD45^+^ cells was not changed (Fig. 6G-H). We also tested the expression of pro-tumor cytokines that secreted by CD8^+^ T cells. qRT-PCR showed Tnfa and Ltβ was decreased significantly with atRA treatment compared to untreated, similar to metformin treatment (Fig. 6I). These data indicated the inhibition of HCC by metformin is at least partly due to the promotion of elevated atRA level.

### scRNA-seq of immune cells of *Fah^-/-^* mouse model reveals hepatic resident-like CD8*^+^* T cells are gradually increased following the development of chronic liver injury

A previous study has reported that the CLI environment of *Fah*^-/-^ mouse model enables to promote the carcinogenesis of implanted HBsAg^+^ hepatocytes and found that HBsAg-specific CD8^+^ T cells participate in this process (35). To figure out whether CD8^+^ T cells could express pro-tumor cytokines and atRA receptor, we obtained related scRNA-seq data. In order to confirm the consistency of transcriptome level with the conclusion we found using flow cytometry assays, CD8^+^ T cells were divided into Cd8a-high expression group and Cd8a-low expression group to better observe the dynamic changes of CD8^+^ T cells during tumor progression at the mRNA level. The distribution of CD8^+^ T cells with high Cd8a expression was more likely to be in the same clusters while CD8^+^ T cells with low Cd8a expression were distributed separately and mixed with other immune cells (Fig. 7A). Interestingly, results showed CD8^+^ T cells will gradually increase after suffering CLI, especially CD8^+^ T cells with high Cd8a expression (Figs. 7A-C and S4A-C). Then, we could find that the cell proportion of CD8^+^ T cells expressing pro-tumor cytokines/molecules was higher than other immune cells, especially Ltb, Pdcd1, Cxcr6 and Il1b (Fig. 7D). And expression levels of most pro-tumor cytokines were higher in CD8^+^ T cells with high Cd8a expression (Fig. S4D). In addition, the expression levels of atRA receptor (Rara and Rarb) were gradually increased following the CLI time (Figs. 7E, S4E). However, CD8^+^ T cells tended to express Rara while little could express Rarb (Figs. 7F, S4F). And the expression level of both Rara and Rarb were correlated with CD8^+^ T cells (Figs. 7G, S4G). Then, we further explored the expression characters of pro-tumor molecules in CD8^+^ T cells. Intriguingly, Tnf, Pdcd1, Cxcr6, Il1b and Il6, excluding Ltb, were all gradually up-regulated along the tumorigenesis of *Fah^-/-^* mouse model (Figs. S5-10A). Then, Tnf, Ltb, Pdcd1, Il1b and Cxcr6, excluding Il6, tended to highly express and distributed in the same CD8^+^ T cell clusters (Figs. S5-10B). And all these pro-tumor genes were correlated with the expression of Cd8a (Figs. S5-10C) and Rara (Figs. S5-10D). In short, CD8^+^ T cells were responsible for the liver tumorigenesis of *Fah*^-/-^ mouse via expressing pro-tumor molecules.

**Figure 7.**
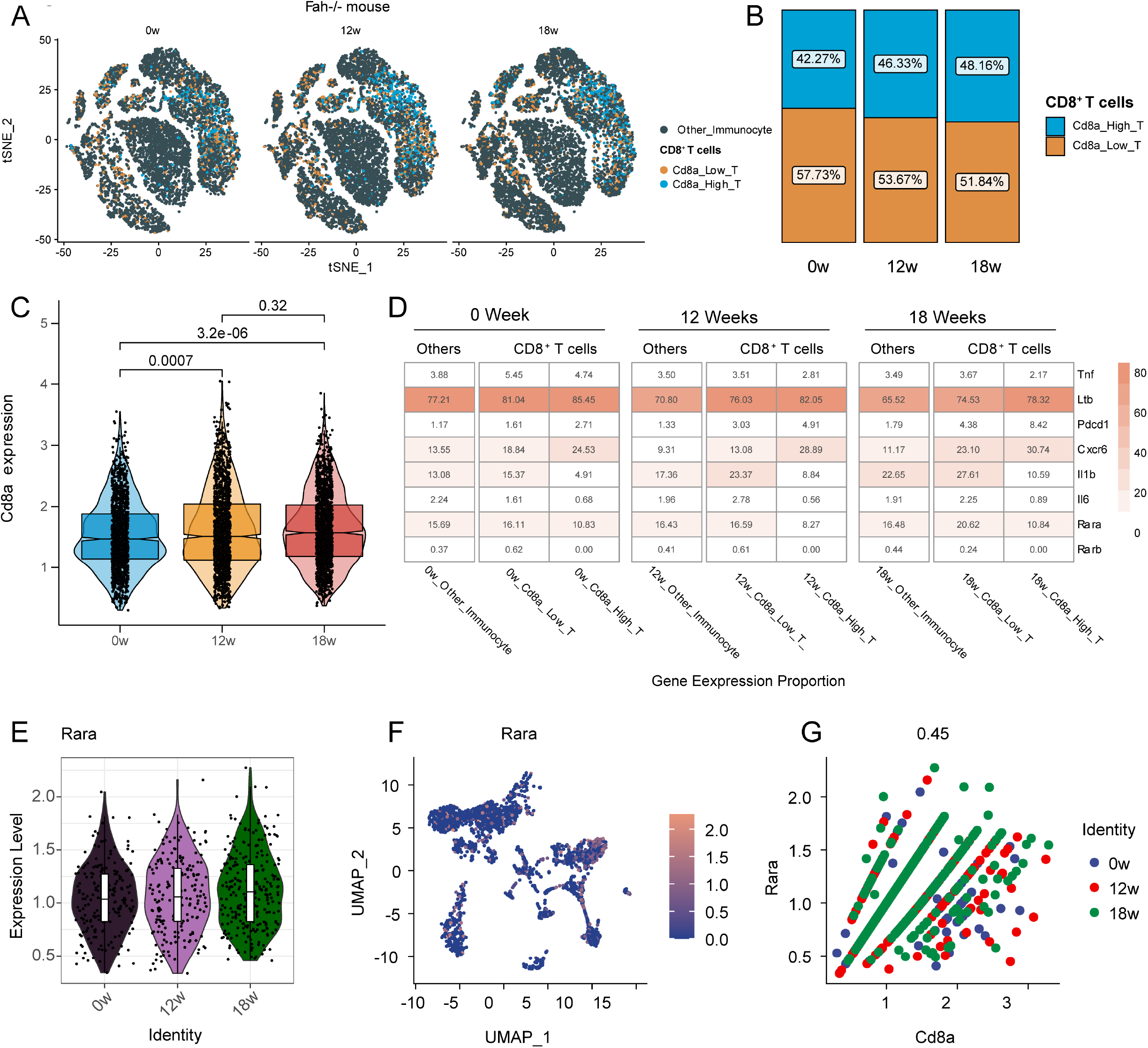
Single cell RNA-sequencing analysis reveals the expression characteristics of CD8*^+^* T cells in *Fah^-/-^* mouse with progressive chronic liver injury. (A) T-distributed stochastic neighbor embedding (t-SNE) plot of immune cells of *Fah^-/-^* mouse at 0, 12, and 18 weeks under CLI. (B) Histogram indicating the proportion of the CD8^+^ T cell at 0, 12, and 18 weeks. (C) Violin plot presenting the expression levels of Cd8a at 0, 12, and 18 weeks. (D) Expression proportion of Tnf, Ltb, Pdcd1, Cxcr6, Il1b, Il6, Rara and Rarb in CD8^+^ T cell and other immune types group at 0, 12, and 18 weeks. (E) Violin plot presenting the expression levels of Rara at 0, 12, and 18 weeks. (F) Uniform manifold approximation and projection (UMAP) plot showing the expression levels of Rara. (G) Correlation analysis of Cd8a and Rara in *Fah^-/-^* mouse at 0 (blue), 12 (red), and 18 weeks (green).

### Hepatic resident-like CD8*^+^* T cells are increased in patients with chronic liver injury

To assess the relevance of our findings between precancerous mouse livers and human HCCs, adjacent liver tissue and HCC samples from The Cancer Genome Atlas (TCGA) and normal liver tissue from GTEx database were downloaded and analyzed for the expression levels of CD8A and pro-tumor cytokines. We found that CD8^+^ T cells increased in the adjacent tumor tissues compared with both HCC tissues and normal liver tissues (Fig. 8A). Also, we compared healthy controls (HC), simple steatosis (SS) and nonalcoholic steatohepatitis (NASH) from GSE89632 database, and found the number of CD8^+^ T cells increased in the human chronic injured liver compared with normal liver (Fig. 8B). Notably, previous paper also found that the fraction of CD8^+^ T cells are higher in HCC and HCC adjacent tissue than in healthy liver tissue, while HCC adjacent tissue contained even more T cells than HCC (58), in concordant with our analysis (Fig. 8A). We also found that all pro-tumor cytokines were up-regulated in adjacent tumor tissues while they were not consistent in tumor tissues when both compared to normal tissues (Figs. 8C and 8D, S11). Moreover, we found a significant higher positive regression coefficient between CD8A expression and TNF or LTβ in adjacent tumor tissues other than HCC tissues and normal tissues (Figs. 8D, S12-14A). In parallel, other four pro-tumor molecules had consistent pattern like TNF or LTβ (Figs. S12-14B). And pan-tissues and pan-cancers analyses show significant correlation between CD8^+^ T cells and these pro-tumor cytokines in most tissues and cancer types, especially liver tissues and HCC, hinting that CD8^+^ T cells contributed to the expression of pro-tumor cytokines (Figs. 8D, S12-14B). Taken together, these results may indicate that the CD8^+^ T cells could express pro-tumor cytokines and might have different expression levels and functions in adjacent tumor tissue and HCC tissues.

**Figure 8.**
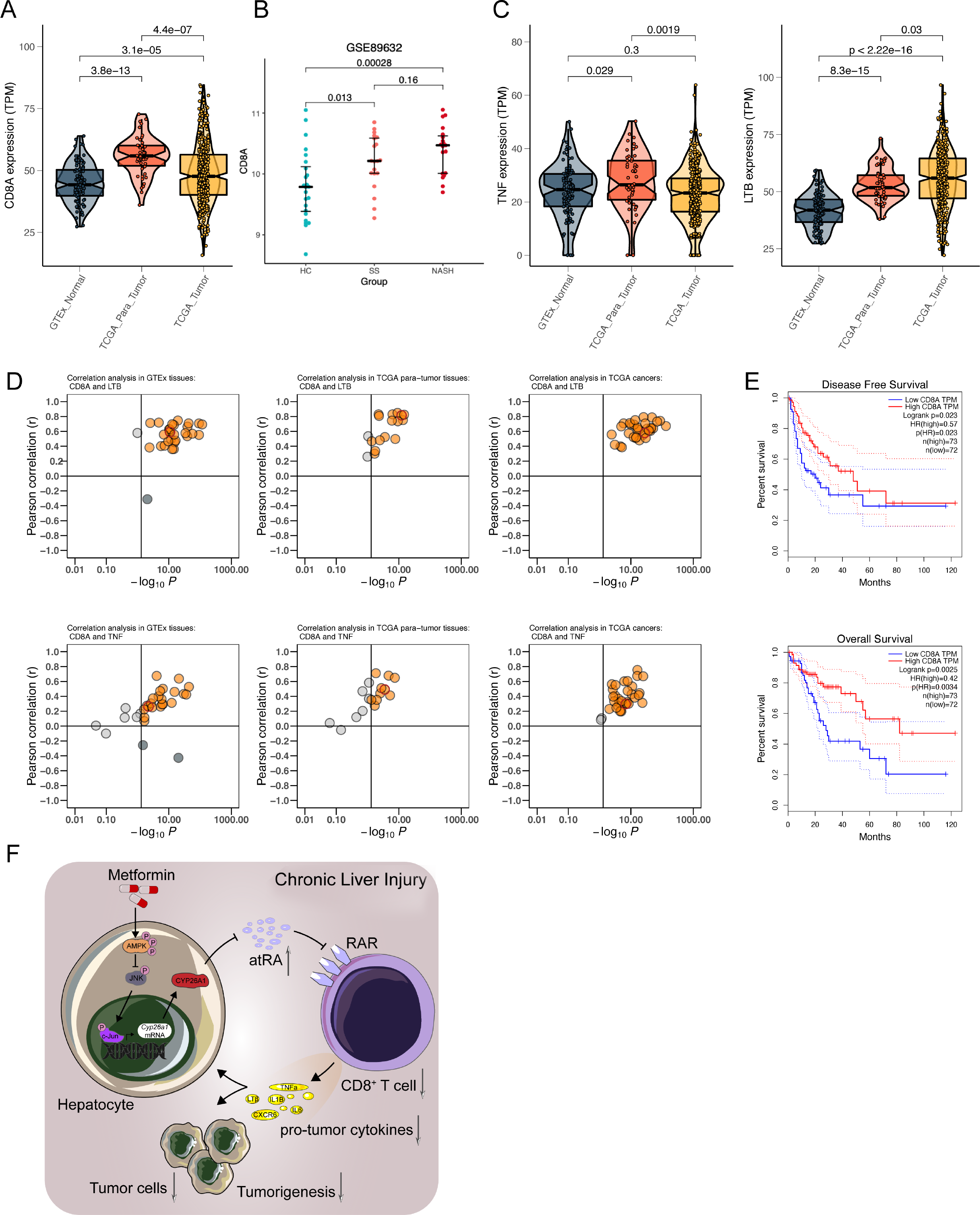
CD8*^+^* T cells have different role in HCC and HCC adjacent tumor based on their molecular and functional characteristics. (A) Violin plot presenting the expression levels of CD8A in GTEx normal, TCGA para-tumor and TCGA tumors tissues. (B) Boxplot presenting the expression levels of CD8A in GSE89632 HC, SS and NASH tissues. (C) Violin plot presenting the expression levels of TNF and LTB in GTEx normal, TCGA para-tumor and TCGA tumors tissues. (D) Correlation analysis of CD8A and LTB or TNF in 30 GTEx normal, 17 TCGA para-tumor and 34 TCGA tumors tissue types. Red color indicates the Liver organ in GTEx or HCC para-tumor tissues, and HCC tumor in TCGA tumor tissues. (E) Kaplan-Meier survival analysis of overall survival (OS) (left) or disease-free survival (DFS) for CD8A transcript level of HCC patients from TCGA database. (F) The schematic diagram illustrating the mechanism of appropriately metformin downregulated tumor promotion in hepatocarcinogenesis. The icons of cell types were obtained from SERVIER MEDICAL ART (https://smart.servier.com/).

Next, we examined the clinical significance of CD8A in liver cancer. When patients were divided into “high” and “low” CD8 expression based on top 20% value of CD8A, we observed that elevated CD8A expression was associated with better patient survival in TCGA liver cancer database (Fig. 8E). This association means CD8^+^ T cells may function as pro-tumor cells in adjacent tumor tissues while as anti-tumor cells in tumor tissues, dependent on the expression levels of pro-tumor or anti-tumor cytokines secreted by CD8^+^ T cells. Altogether, these results showed the number and function of hepatic resident-like CD8^+^ T cells were changed in livers with varying degrees of injury (normal liver, chronic injured liver and HCC).

Taken together, these results show metformin suppressed the expression of *Cyp26a1* through the AMPK/JNK/c-Jun pathway and leading to the elevation of atRA level. The atRA treatment inhibited CD8^+^ T cells proliferation by binding to atRA receptor (RAR) on the cell membrane of CD8^+^ T cells. The decrease of pro-tumor CD8^+^ T cells can reduce the release of pro-tumor cytokines (especially Tnfa and Ltβ) and inhibit hepatocarcinogenesis (Fig. 8F)

## Discussion

Currently, systemic options for the prevention and treatment of HCC are limited. New therapeutic targets, preventive strategies as well as biomarkers for patient stratification need to be discovered. Therefore, it is a wise choice to test new therapeutic targets and preventive strategies using animal model. HCC development is a complex and multistep process caused by diverse risk factors, determining that no adaptive animal model can fully mimic the occurrence and development of human HCC. Human *Fah^-/-^* induced HCC occurs in a very low incidence. The murine model of *Fah*-deficiency is a suitable animal model, which represents all phenotypic and biochemical characterizations of patients with FAH deficiency (24). Furthermore, the HCC model of CLI *Fah^-/-^* mice is highly similar to human alcohol-induced HCC or c-myc-altered HCC (25). So far, there are many studies using this animal model to discover and verify relevant biological phenomena and mechanisms related to HCC. These indicates that the study of *Fah*-deficient chronic injury mice could be a good indicator of human HCC.

In this study, we revealed altered transcriptomic profiles and signaling pathways in the early stages of chronic liver injury during hepatocarcinogenesis, and identified hepatic responses to metformin treatment. Our results uncovered that the inhibition of AMPK activity and the symptoms of hypoglycemia are co-existent in the model of chronic liver injury (Fig. 1D, F). Besides, AMPK activity is repressed in kinds of human liver disease (Fig, 1G-I) (37, 38). AMPK activity is reduced by inflammation, obesity, and diabetes. The activation of AMPK pathway has been viewed as a viable therapeutic strategy to improve HCC (59). These results suggest that metformin may improve various liver diseases by up-regulating the repressed AMPK activity under disease conditions.

Metformin is an AMPK activator and blood glucose regulator, which prompted us to investigate the role and mechanism of metformin in our mouse HCC model. Epidemiological studies have shown a reduction in incidence rate and mortality of liver cancer in type 2 diabetes patients treated with metformin (60–62). In our mouse model, metformin treatment significantly reduced the HCC incidence of *Fah^-/-^* mice (Fig. 2C-E). Chronic inflammation contributes to hepatocarcinogenesis through multiple mechanisms, one of which is the secretion of pro-tumor cytokines by immune cells. The number and proportion of CD8^+^ T cells increased significantly during chronic liver injury, and metformin could reverse this effect in *Fah^-/-^* mice (Fig. 6B-D). CD8^+^ T cells could be classified based on their molecular and functional characteristics. Generally, tumor infiltrating CD8^+^ T cells are anti-tumor immune cells and associated with a favorable prognosis (63, 64). On the other hand, CD8^+^ T cells with high expression of inhibitory receptors (PD-1, TIGIT, TIM-3, LAG-3) is defined as exhausted CD8^+^ T cells, which progressively loses effector function and possesses a poor outcome (65, 66).

Recently, several studies showed CD8^+^ T cells contributes to HCC development in many types of mouse chronic liver injury (19, 25, 56, 57). Endig *et al.* showed that CD8^+^ T cells and LTβ signaling contributes to HCC development in Fah^-/-^ mice with chronic liver injury, which is exactly the same as the mouse model we used, but the way of inducing chronic liver injury is slightly different (25). Pfister *et al.* reported in a preclinical model of NASH-induced HCC, CD8^+^ T cells contribute to the induction of NASH-HCC rather than invigorating or executing immune surveillance (57). Concordantly, another group has revealed that auto-aggression of CD8^+^ T cells in the liver may be involved in the development of HCC in patients with NASH (56). Williams *et al.* also observed the enrichment of exhausted CD8^+^ T cells in the precancerous liver of *Ncoa5*^+/−^ mouse and these T cells function as a pro-tumor microenvironment. Meanwhile, metformin treatment is able to reduce hepatic infiltration of CD8^+^ T cells and reduce HCC formation in the chronic liver injury of *Ncoa5*^+/−^ mice(19). All these findings suggest that the classification (effector or exhaustion) and location (tumor or adjacent tumor) of CD8^+^ T cells determine whether they are pro-tumor or anti-tumor T cells. Our data proved the number and ratio of CD8^+^ T cells were increased in the process of chronic liver injury, and metformin could reduce the number and ratio of CD8^+^ T cells in liver and then suppressed the HCC formation in *Fah^-/-^* mice (Figs. 6B-D). The single cell RNA-seq analysis of CD8^+^ T cells in *Fah^-/-^* mice reveals that CD8^+^ T cells could express pro-tumor cytokines and atRA receptors (Fig. 7). Interestingly, this study also found that the fraction of CD8^+^ T cells was higher in HCC and HCC adjacent tissues than in healthy liver tissue, while HCC adjacent tissue contained even more CD8^+^ T cells than HCC (Fig. 8) (58). Thus, in addition to gaining insights into the mechanism of actions of metformin, our data could be valuable in supporting that metformin treatment may be beneficial in reversing CD8^+^ T cell-exhausted tumor microenvironment for HCC patients.

Retinoic acid (RA) is an active metabolite of vitamin A. Retinoic acid includes all-*trans* retinoic acid (atRA), 9-*cis*-retinoic acid (9-*cis* RA), and 13-*cis*-retinoic acid (13-*cis* RA)(41). Vitamin A is irreversibly converted to retinoic acid or atRA by the aldehyde dehydrogenase (ALDH) enzyme family members. The atRA was degraded by atRA-degrading cytochrome P450 reductases, such as Cyp26a1, which converts atRA to inactive metabolites(41). The up-regulation of ALDH or down-regulation of Cyp26a1 will lead to the increase of atRA level. In our study, we found the expression of Cyp26a1 was decreased by metformin treatment *in vivo* and *in vitro* (Fig. 3). The decrease of *Cyp26a1* gene expression will result in the increase of atRA level. Besides, metformin directly increased the luciferase activity of atRA response element plasmid, further demonstrating that metformin promotes the level of atRA in hepatocytes (Fig. 3J). Preclinical and clinical studies of atRA against tumor are reported increasingly. A classic example of clinical use is the treatment of acute promyelocytic leukemia (APL), which is the most efficacious use of atRA in cancer therapy (42, 43). Bryan *et al.* investigated that atRA combined with paclitaxel had better overall clinical efficacy than paclitaxel alone in the treatment of recurrent or metastatic breast cancer (44). Han *et al.* showed that the storage of retinaldehyde was significantly decreased in patients with HCC and found that retinol metabolism has great prospects for clinical application in diagnosis, prognosis and chemotherapy of HCC (67). There are several mechanisms for the anti-tumor activity of atRA, one of which is the effect on the immune systems (45, 68). In our study, we also proved that atRA agents possess anti-tumor activity through their ability to reduce the number and ratio of CD8^+^ T cells in *Fah^-/-^* mice (Fig. 6). CD8^+^ T cells expressed the atRA receptor RAR (Fig. 7 and S4E) and responded to atRA (69). The atRA treatment inhibited T cell proliferation in a dose-dependent manner *in vivo* and *in vitro* (68, 70). These may illustrate metformin can reduce the number and ratio of CD8^+^ T cells through elevate the level of atRA in our mouse model. Metformin/Cyp26a1/atRA signaling pathway may partly explain how metformin affected CD8^+^ T cell proliferation and differentiation.

Metformin and atRA are both identified as anti-neoplastic agents. Our data show both metformin and atRA reduce the number and proportion of CD8^+^ T cells which was considered as pro-tumor immune cells in our mouse model (Fig. 6). In summary, we firstly revealed that metformin increases the level of atRA by inhibiting the expression of *Cyp26a1* gene, and then inhibits HCC caused by chronic liver injury (Fig. 8F).

## Acknowledgments

The authors thank Yanxiang Song, Hua Qiu, Cheng Wang for tissue preparation, sectioning, and staining; Xiuhua Li, Changcheng Liu for Flow Cytometric analysis; Yanli Lu, Xiaolan Mu, Xinyue Du, Shengwei Shen, Mingyang Xu, Shihan Sun for their kindly assistance and meaningful discussion during manuscript preparation.

This work was funded by Major Program of National Key Research and Development Project (2020YFA0112600, 2019YFA0801502), National Natural Science Foundation of China (82002945, 82173019, 82103095), Jiangxi Provincial Natural Science Foundation (20212ACB206033), the Project of Shanghai science and technology commission (19140902900), Program of Shanghai Academic/Technology Research Leader (20XD1434000), and Peak Disciplines (Type IV) of Institutions of Higher Learning in Shanghai.

## Author Contributions

WH, MC, CL designed and performed experiments, analyzed data; WC, YY, YC, ZY performed animal experiments; XW, GW, LP performed bioinformatics analysis; WH, YC performed immunohistochemistry analysis; ZH, JW, QT supervised and planned research; ZH conceptualized study, supervised and planned research. WH, ZH wrote the paper.

## Data Availability Statement

All data generated or analysed during this study are included in the manuscript and supporting file; Source Data files have been provided for Figures 1 and 3.

---

**Figure S1.**
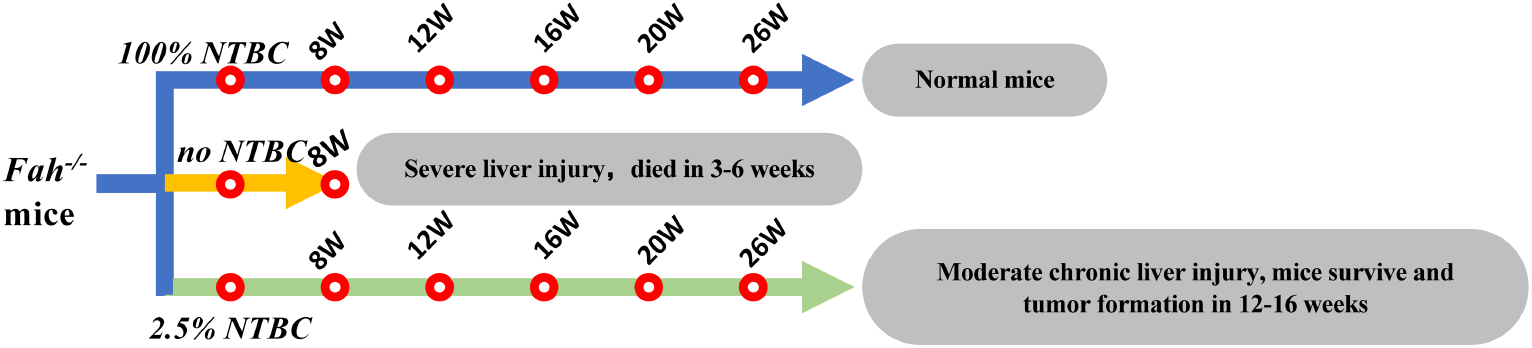
characteristics of *Fah*^-/-^ mice. under 100% NTBC, the liver of Fah^-/-^ mice was preserved normally; Fah^-/-^ mice developed acute liver injury and died at 3-6 weeks without any NTBC; Fah^-/-^ mouse can survive under 2.5% NTBC, but experience chronic liver injury and form liver cancer after 12 weeks.

**Figure S2.**
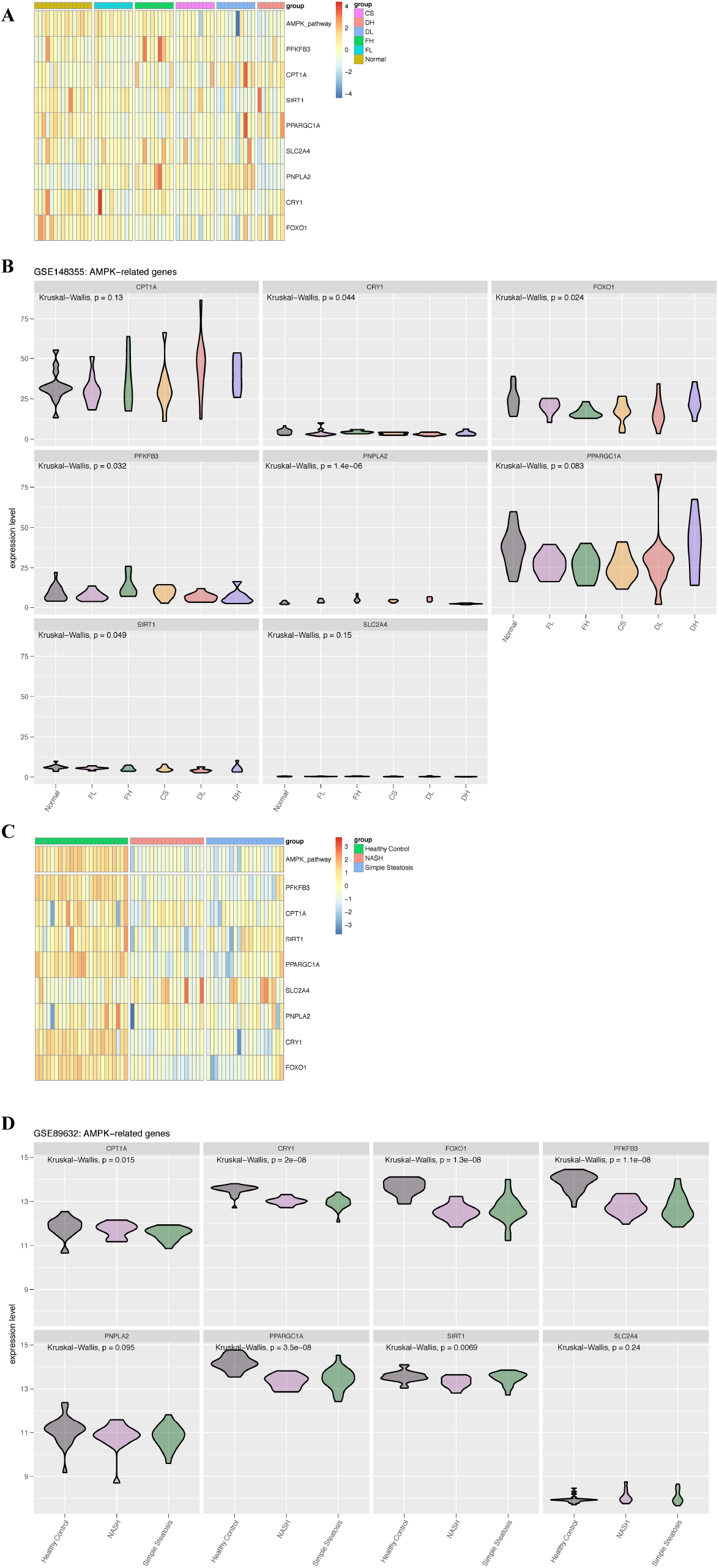
AMPK pathway was suppressed in chronic liver injury tissues compare with normal tissues. (A)Heatmap of AMPK pathway and related genes in normal and premalignant tissues. (B)Expression level of AMPK pathway related genes in normal and premalignant tissues (GSE148355 dataset). (C) Heatmap of AMPK pathway and related genes in normal tissues and NAFLD tissues. (D) Expression level of AMPK pathway related genes in normal tissues and NAFLD tissues (GSE89632 dataset). Normal: non-tumor Normal control, FL: low Fibrosis, FH: high Fibrosis, CS: Cirrhosis (CS), DL: Dysplastic nodule Low, DH: Dysplastic Nodule high, NASH: nonalcoholic steatohepatitis

**Figure S3.**
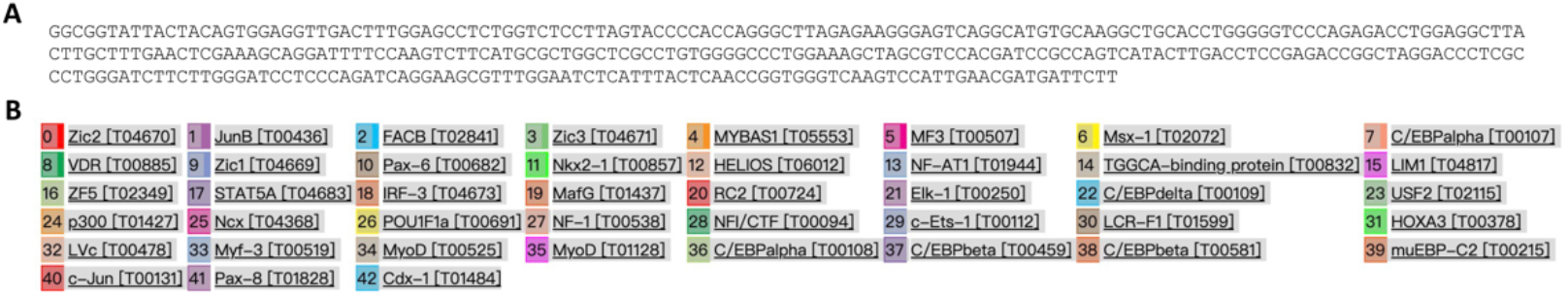
Predict transcription factors that bind in the promoter region of nucleotide – 750 to -500 relation to transcription start site of mouse cyp26a1 gene. (A)The detail sequence of nucleotide -750 to -500 relation to transcription start site of mouse cyp26a1 gene. (B) All the predicted transcription factors binding to nucleotide -750 to -500 relation to transcription start site of mouse cyp26a1 gene using PROMO database (http://alggen.lsi.upc.es/cgibin/promo_v3/promo/promoinit.cgi?dirDB=TF_8.3)

**Figure S4.**
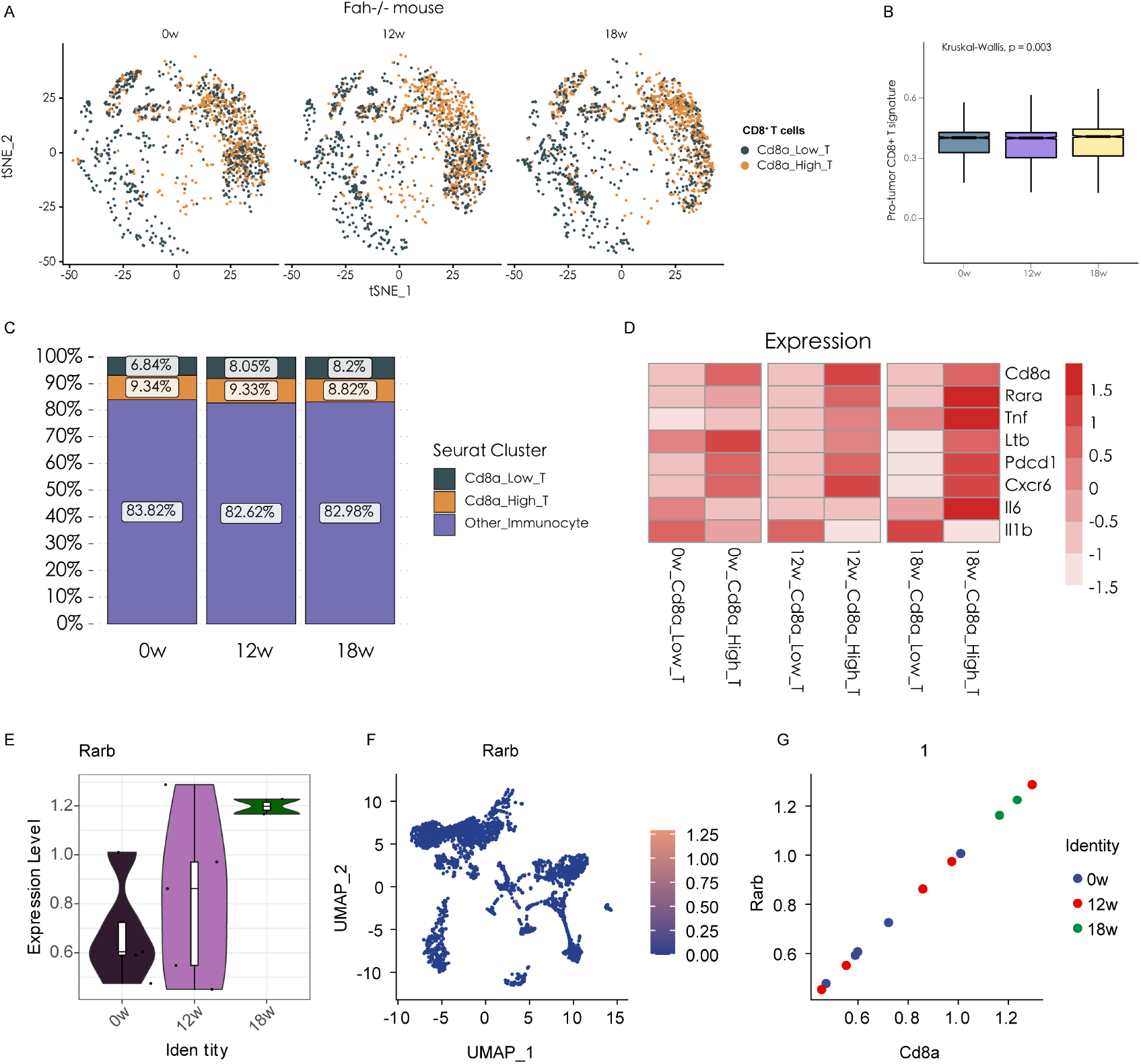
The expression levels of CD8^+^ T cells in *Fah^-/-^* mouse at 0, 12, and 18 weeks. (A) t-SNE visualization of Cd8^High^ T cell (grey) and Cd8^Low^ T cell group (orange) in *Fah^-/-^* mouse at 0, 12, and 18 weeks. (B) Expression levels of tumor promoting CD8^+^ T cell signature in *Fah^-/-^* mouse at 0, 12, and 18 weeks. (C) Histogram indicating the proportion of cells in the Cd8^High^ T cell (grey), Cd8^Low^ T cell (orange) and other immune types group (pruple) at 0, 12, and 18 weeks. (D) Heatmap presenting the expression levels of Cd8a, Rara, Tnf, Ltb, Pdcd1, Cxcr6, Il6 and Il1b in the Cd8^High^ T cell and Cd8^Low^ T cell group at 0, 12, and 18 weeks. (E) Violin plot presenting the expression levels of Rarb at 0, 12, and 18 weeks. (F) UMAP plot showing the expression levels of Rarb. (G) Correlation analysis of Cd8a and Rarb in *Fah^-/-^* mouse at 0 (blue), 12 (red), and 18 weeks (green).

**Figure S5.**
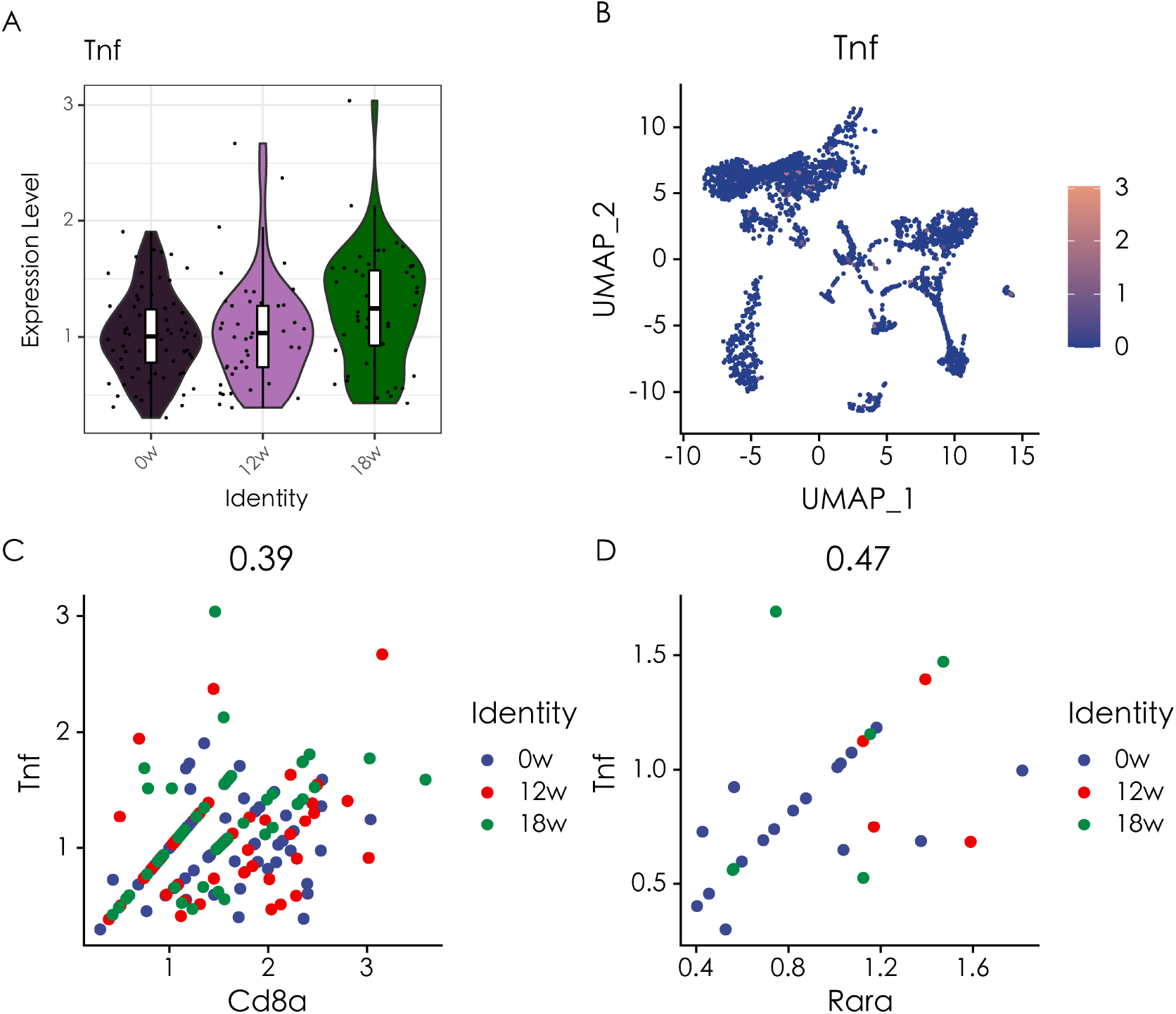
The expression levels of Tnf and its correlation with Cd8a and Rara. (A) Violin plot presenting the expression levels of Tnf at 0, 12, and 18 weeks. (B) UMAP plot showing the expression levels of Tnf. (C) Correlation analysis of Cd8a and Tnf in *Fah^-/-^* mouse at 0 (blue), 12 (red), and 18 weeks (green). (D) Correlation analysis of Rara and Tnf in *Fah^-/-^* mouse at 0 (blue), 12 (red), and 18 weeks (green).

**Figure S6.**
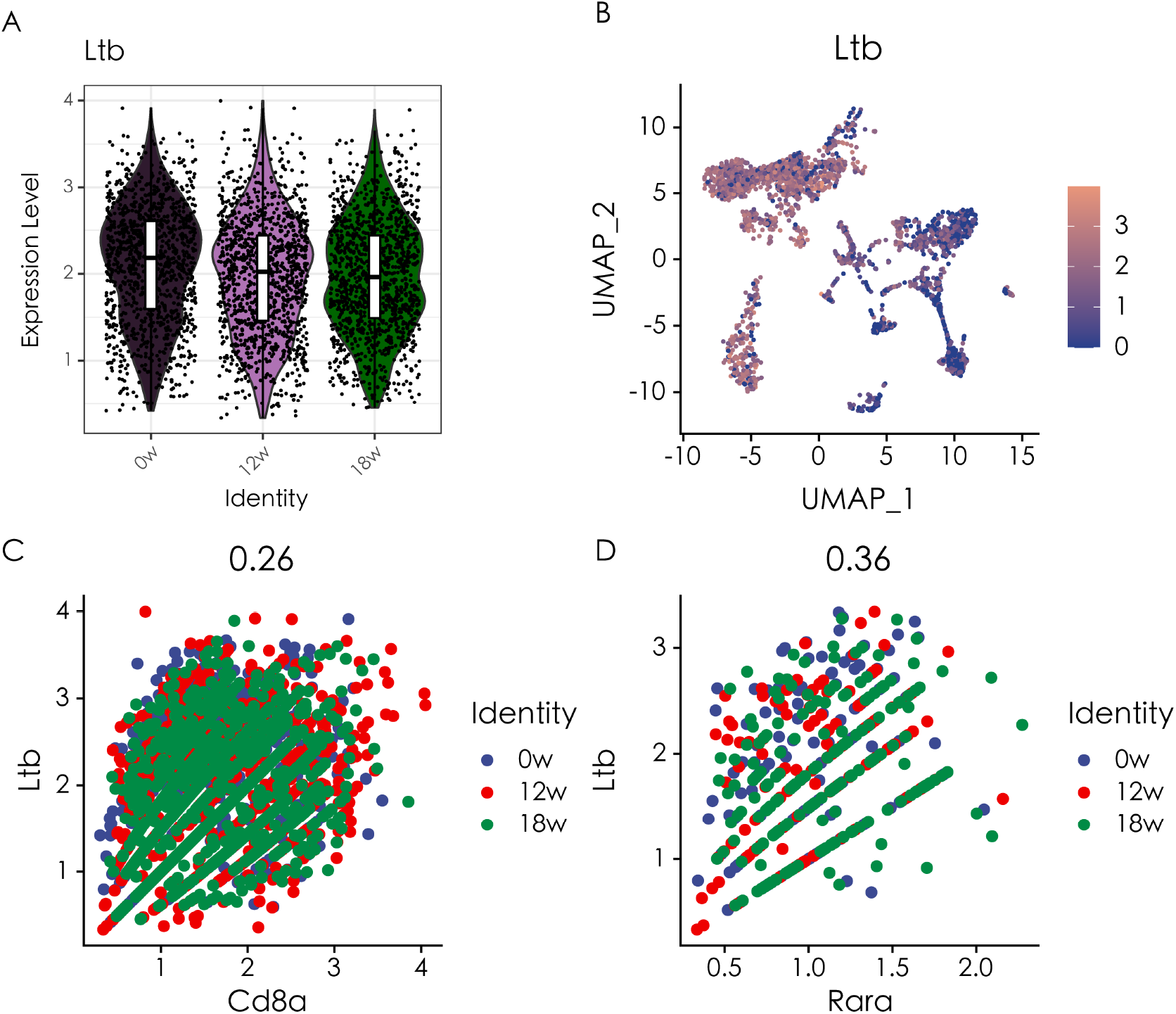
The expression levels of Ltb and its correlation with Cd8a and Rara. (A) Violin plot presenting the expression levels of Ltb at 0, 12, and 18 weeks. (B) UMAP plot showing the expression levels of Ltb. (C) Correlation analysis of Cd8a and Ltb in *Fah^-/-^* mouse at 0 (blue), 12 (red), and 18 weeks (green). (D) Correlation analysis of Rara and Ltb in *Fah^-/-^* mouse at 0 (blue), 12 (red), and 18 weeks (green).

**Figure S7.**
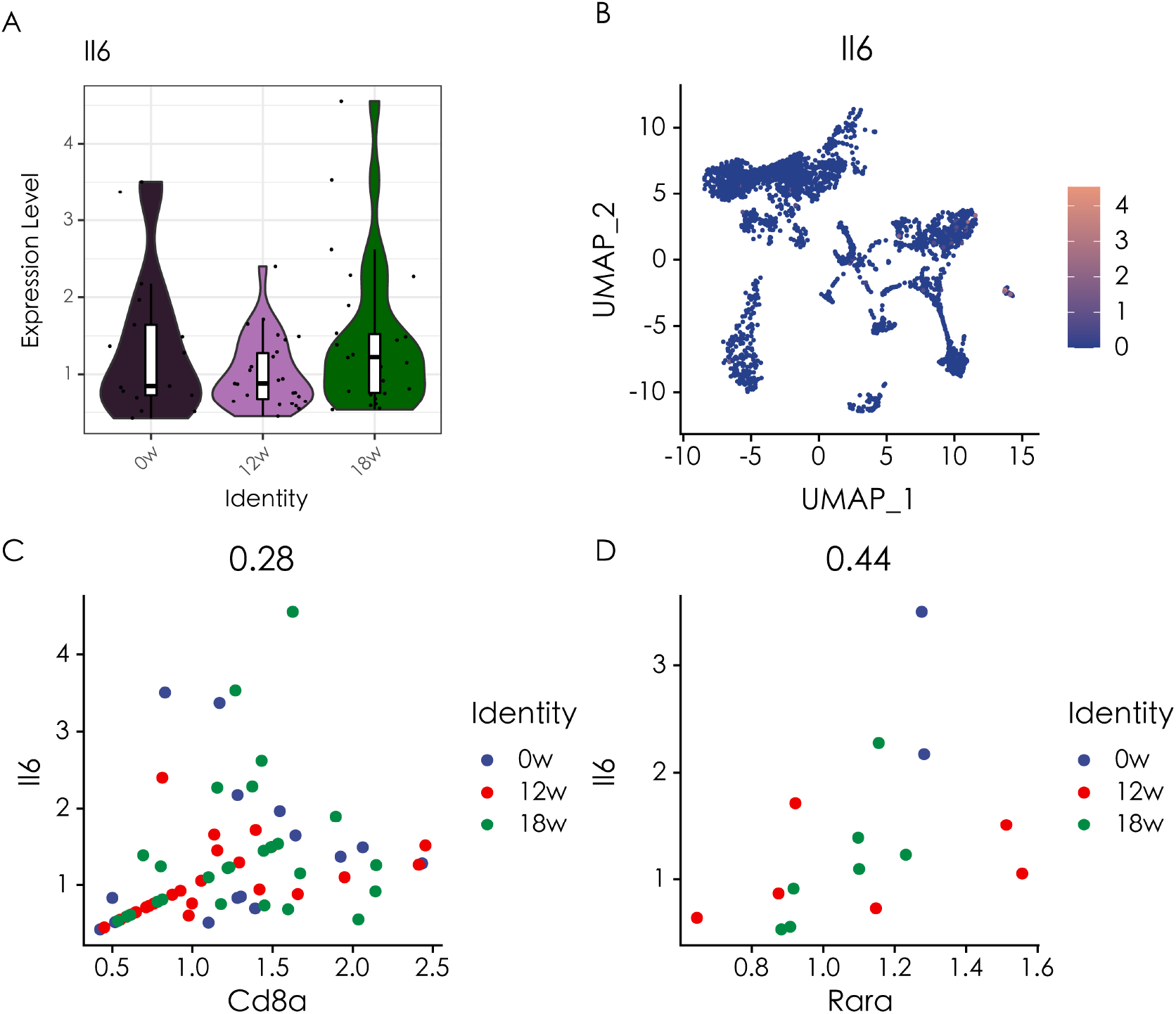
The expression levels of Il6 and its correlation with Cd8a and Rara. (A) Violin plot presenting the expression levels of Il6 at 0, 12, and 18 weeks. (B) UMAP plot showing the expression levels of Il6. (C) Correlation analysis of Cd8a and Il6 in *Fah^-/-^* mouse at 0 (blue), 12 (red), and 18 weeks (green). (D) Correlation analysis of Rara and Il6 in *Fah^-/-^* mouse at 0 (blue), 12 (red), and 18 weeks (green).

**Figure S8.**
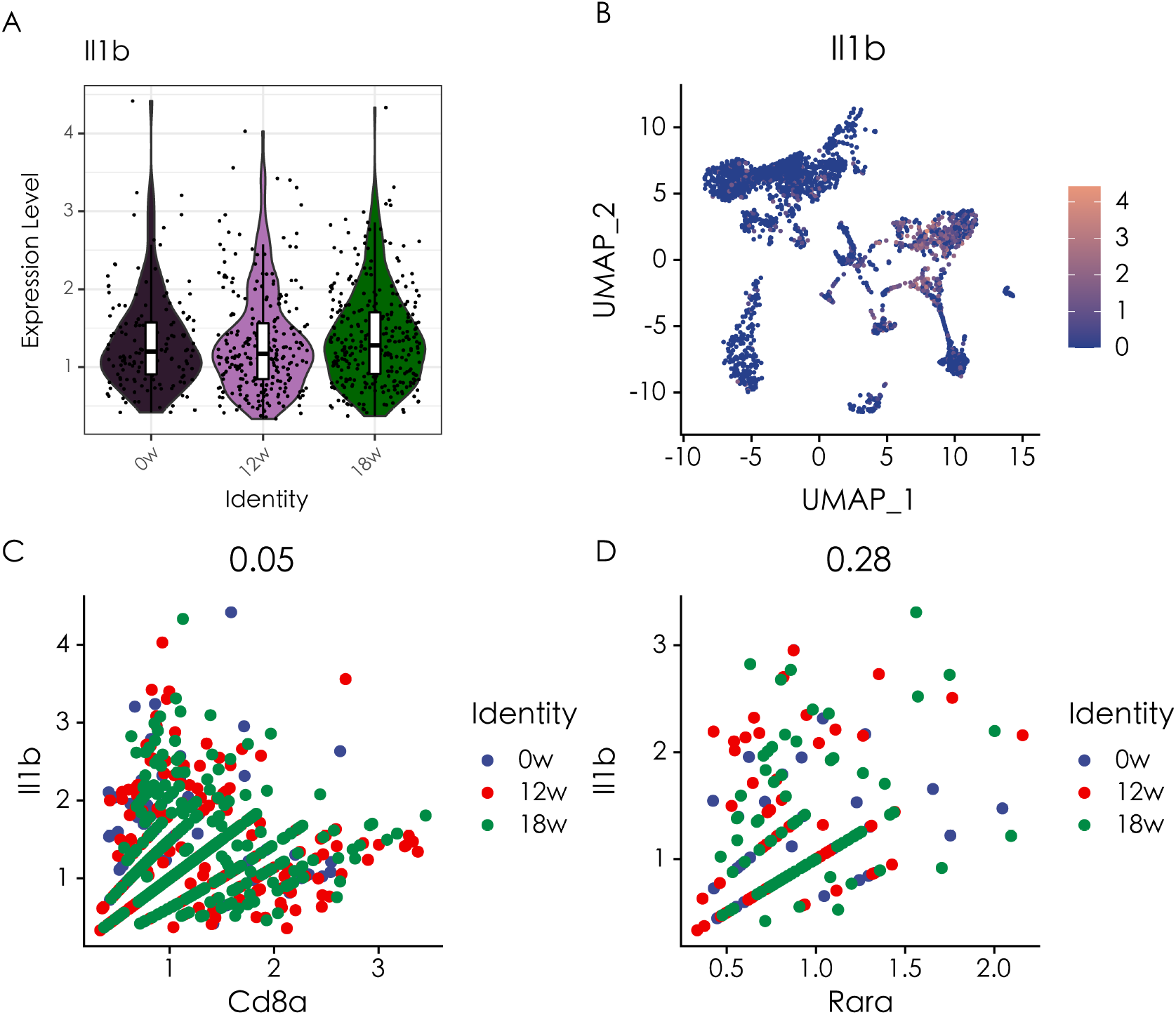
The expression levels of Il1b and its correlation with Cd8a and Rara. (A) Violin plot presenting the expression levels of Il1b at 0, 12, and 18 weeks. (B) UMAP plot showing the expression levels of Il1b. (C) Correlation analysis of Cd8a and Il1b in *Fah^-/-^* mouse at 0 (blue), 12 (red), and 18 weeks (green). (D) Correlation analysis of Rara and Il1b in *Fah^-/-^* mouse at 0 (blue), 12 (red), and 18 weeks (green).

**Figure S9.**
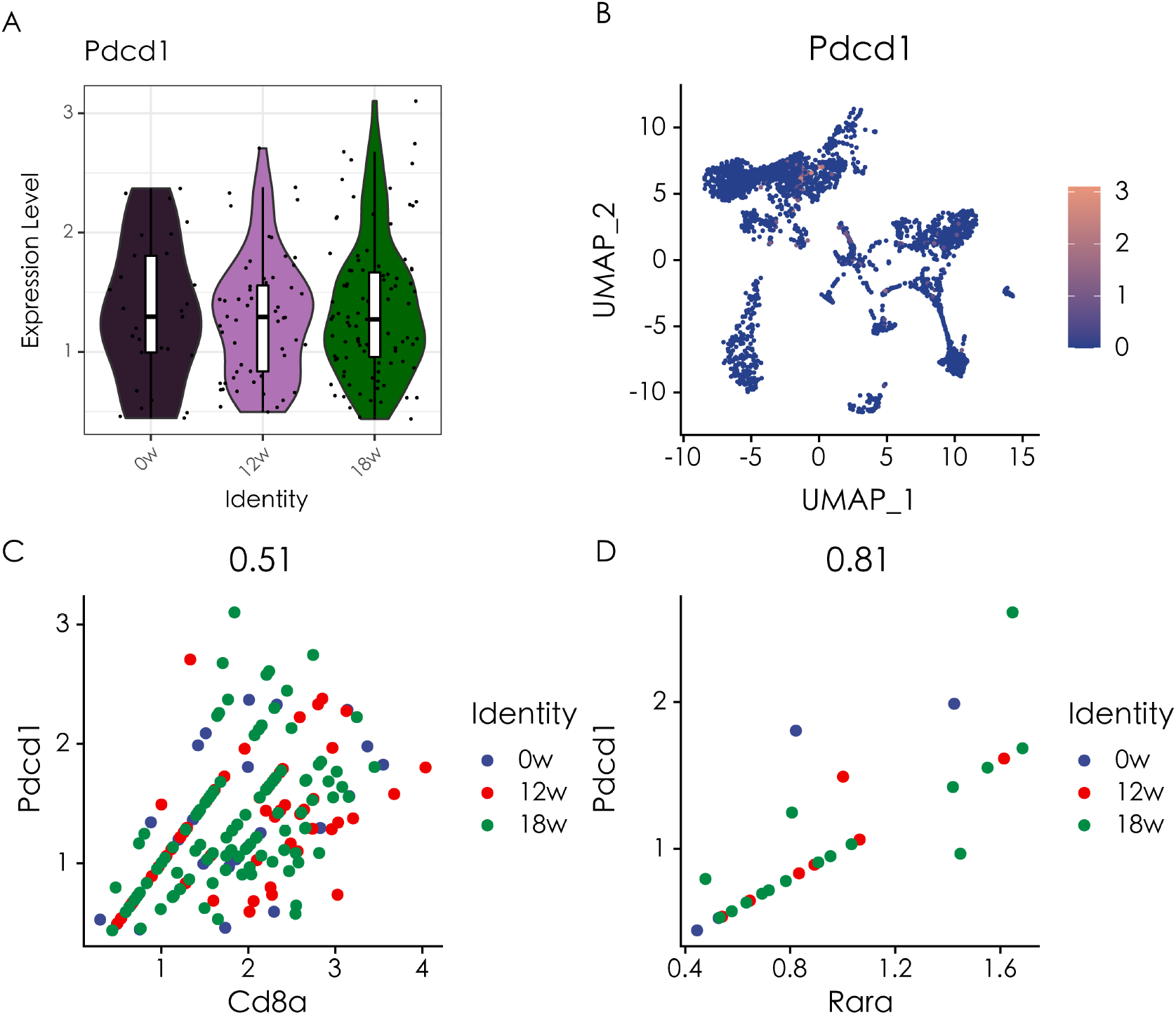
The expression levels of Pdcd1 and its correlation with Cd8a and Rara. (A) Violin plot presenting the expression levels of Pdcd1 at 0, 12, and 18 weeks. (B) UMAP plot showing the expression levels of Pdcd1. (C) Correlation analysis of Cd8a and Pdcd1 in *Fah^-/-^* mouse at 0 (blue), 12 (red), and 18 weeks (green). (D) Correlation analysis of Rara and Pdcd1 in *Fah^-/-^* mouse at 0 (blue), 12 (red), and 18 weeks (green).

**Figure S10.**
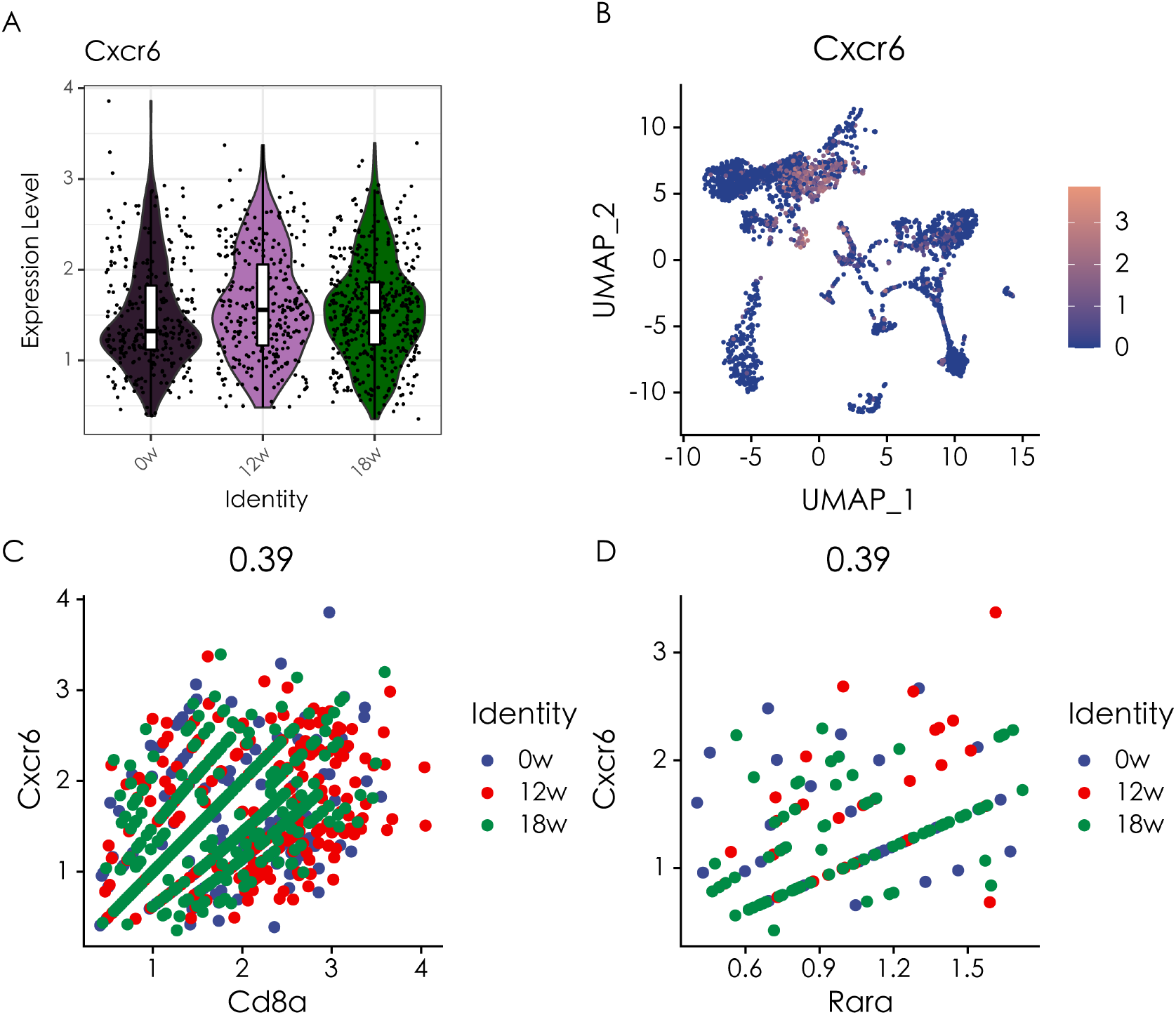
The expression levels of Cxcr6 and its correlation with Cd8a and Rara. (A) Violin plot presenting the expression levels of Cxcr6 at 0, 12, and 18 weeks. (B) UMAP plot showing the expression levels of Cxcr6. (C) Correlation analysis of Cd8a and Cxcr6 in *Fah^-/-^* mouse at 0 (blue), 12 (red), and 18 weeks (green). (D) Correlation analysis of Rara and Cxcr6 in *Fah^-/-^* mouse at 0 (blue), 12 (red), and 18 weeks (green).

**Figure S11.**
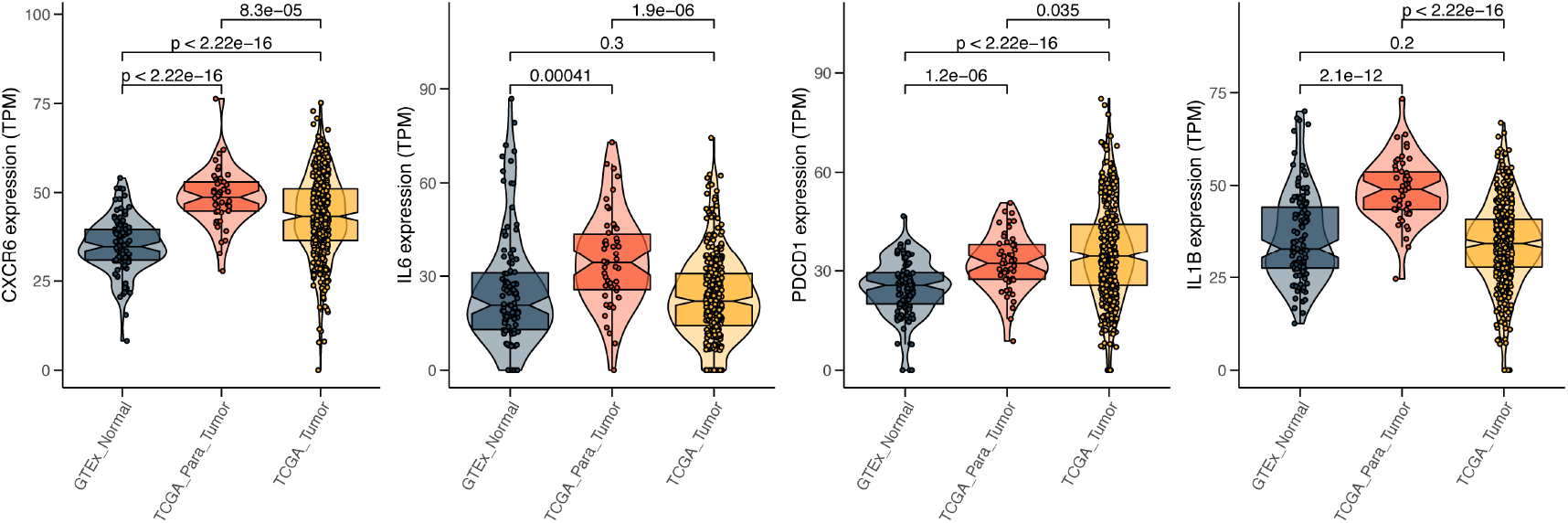
The expression characteristics of CD8+ T cells in GTEx normal, TCGA para-tumor and TCGA tumors tissues. Violin plot presenting the expression levels of CXCR6, IL6, PDCD1 and IL1B in GTEx normal, TCGA para-tumor and TCGA tumors tissues.

**Figure S12.**
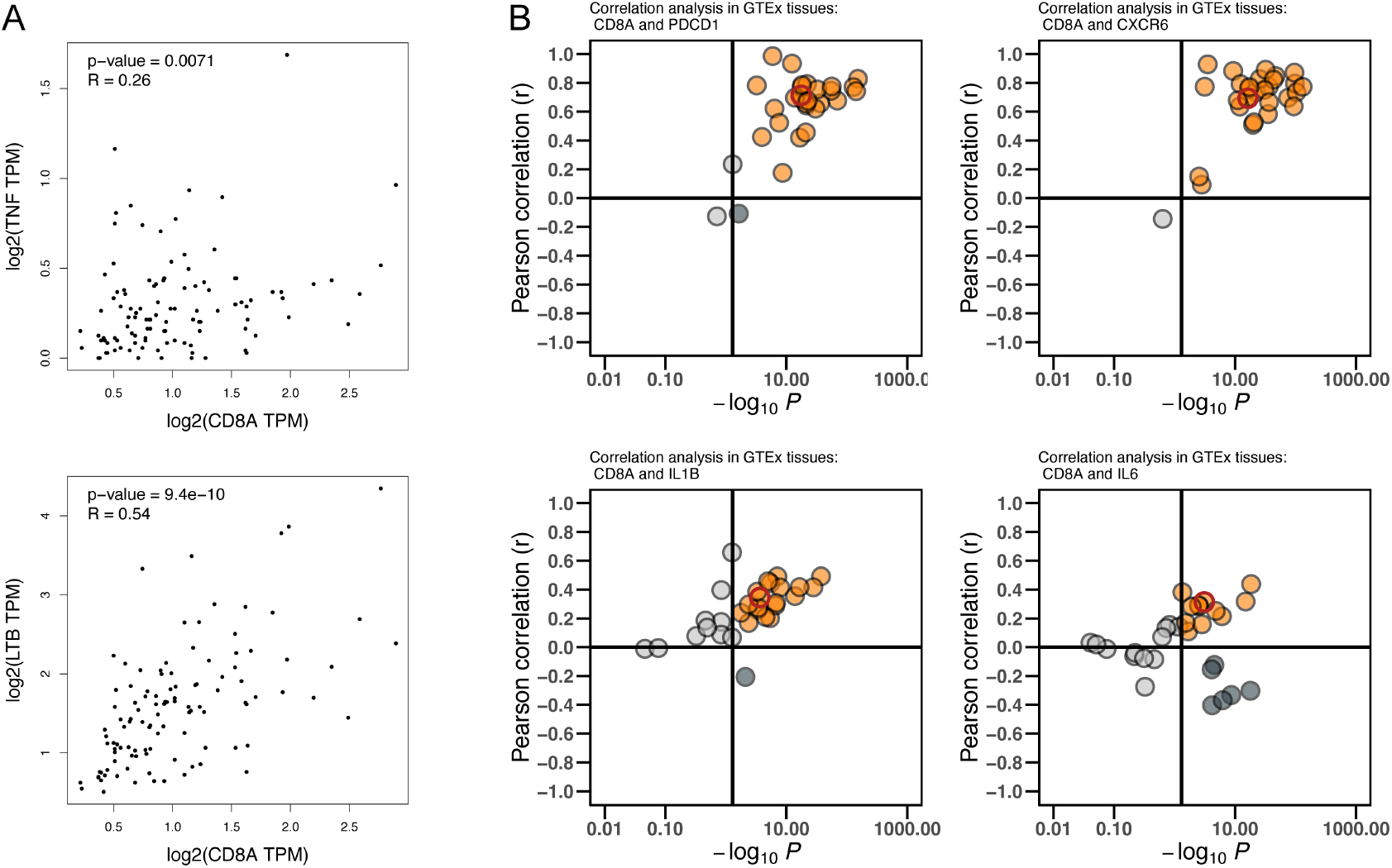
CD8A is positively correlated with pro-tumor cytokines in GTEx normal tissues. (A) Correlation analysis of CD8A and TNF and LTB in 30 GTEx normal tissues. (B) Correlation analysis of CD8A and PDCD1, CXCR6, IL6, IL1B in 30 GTEx normal tissues.

**Figure S13.**
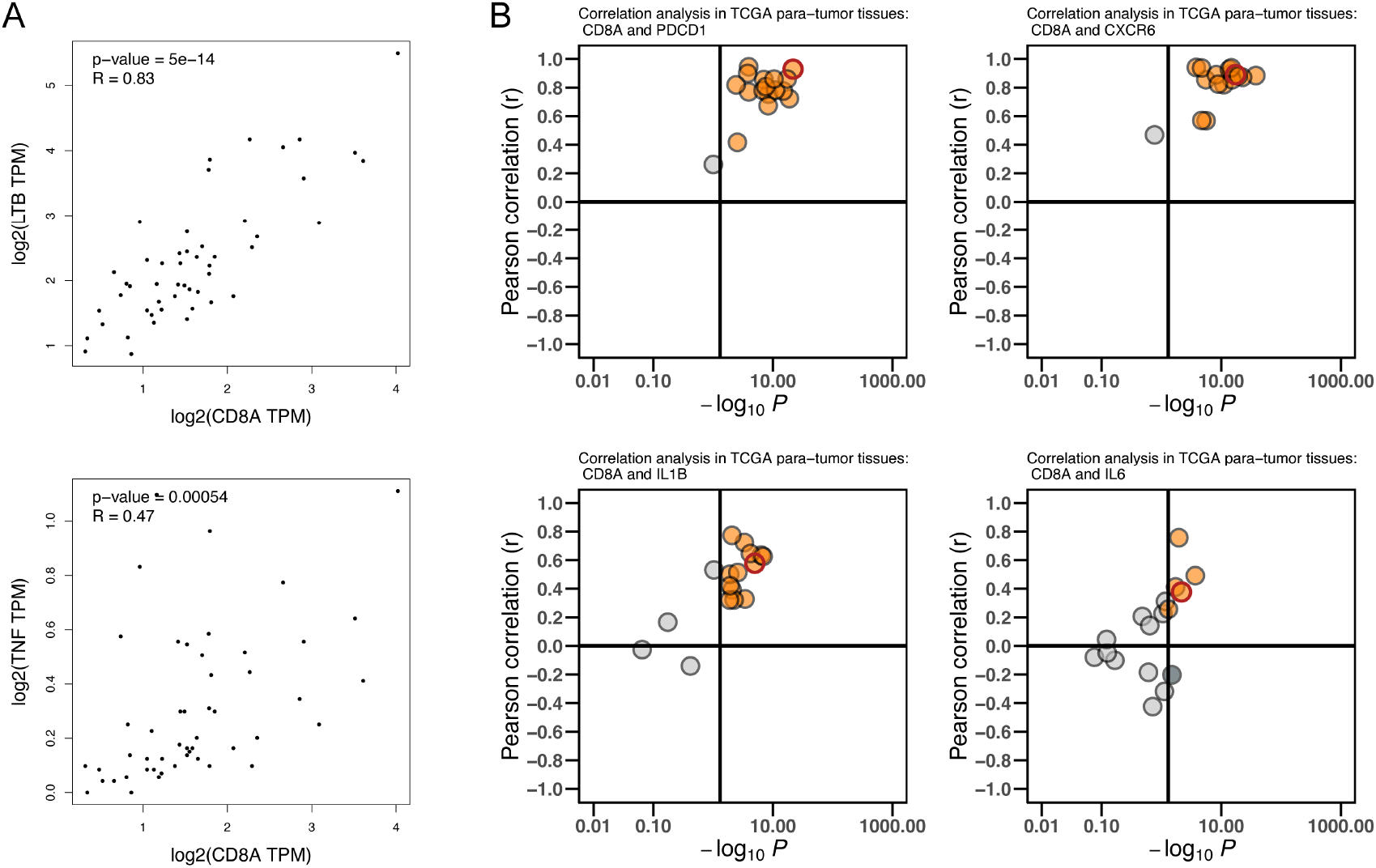
CD8A is positively correlated with pro-tumor cytokines in TCGA para-tumor tissues. (A) Correlation analysis of CD8A and TNF and LTB in 17 TCGA para-tumor tissues. (B) Correlation analysis of CD8A and PDCD1, CXCR6, IL6, IL1B in 17 TCGA para-tumor tissues.

**Figure S14.**
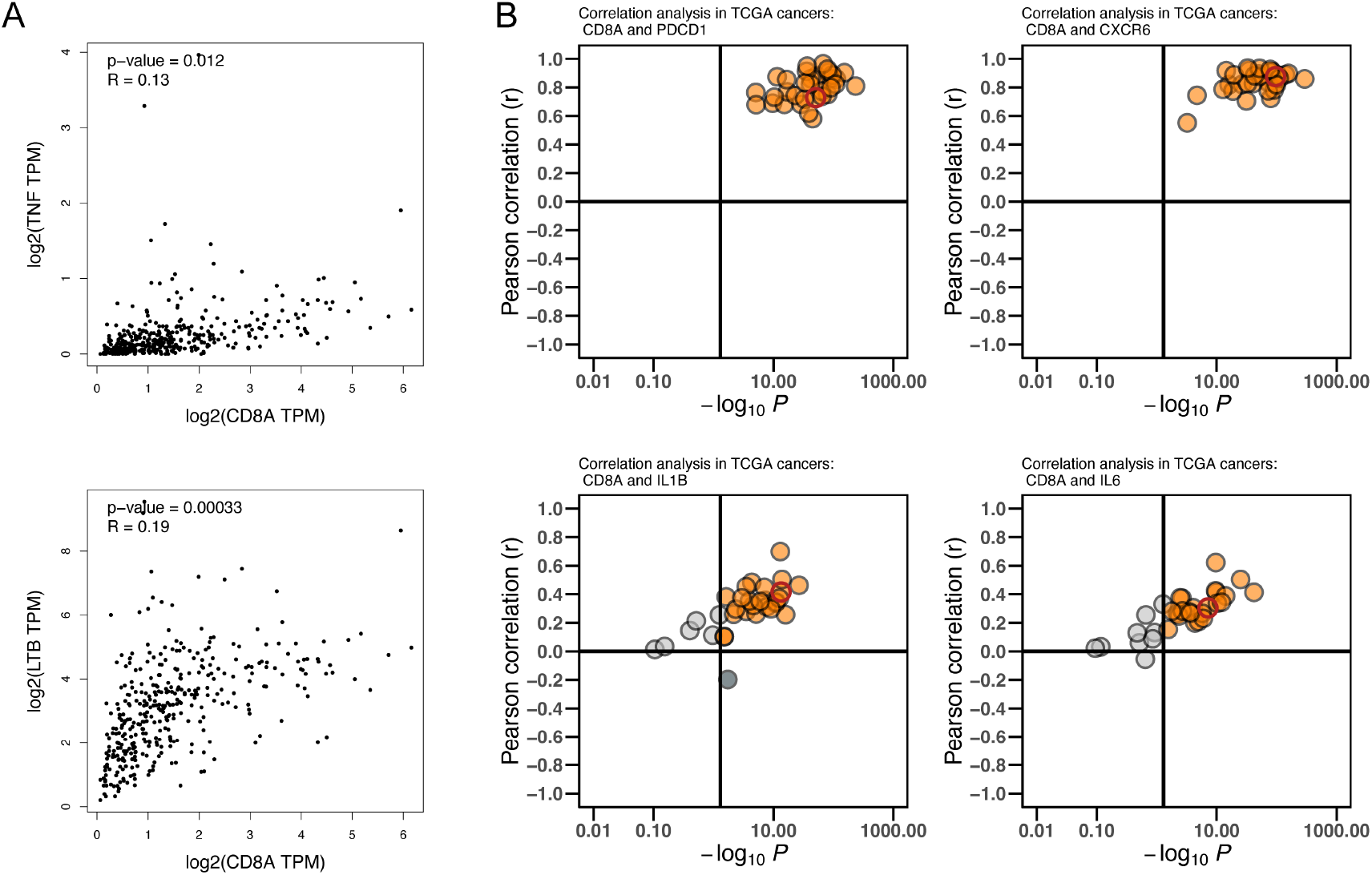
CD8A is positively correlated with pro-tumor cytokines in TCGA tumors tissues. (A) Correlation analysis of CD8A and TNF and LTB in 34 TCGA tumors tissues. (B) Correlation analysis of CD8A and PDCD1, CXCR6, IL6, IL1B in 34 TCGA tumors tissues.

